# Discovery of an AIM2 inflammasome inhibitor for the treatment of DNA-driven inflammatory disease

**DOI:** 10.1101/2022.07.28.501942

**Authors:** Jack P. Green, Lina Y. El-Sharkawy, Stefan Roth, Jie Zhu, Jiayu Cao, Andrew G. Leach, Arthur Liesz, Sally Freeman, David Brough

## Abstract

Inflammation driven by DNA sensors is now understood to be central to disease pathogenesis. Here we describe new inhibitors of pathogenic DNA sensing, primarily of the inflammasome forming sensor AIM2. Molecular modelling and biochemistry has revealed potent inhibitors of AIM2 that work by binding competitively to the DNA binding site. Though less potent, these AIM2 inhibitors, 4-sulfonic calixarenes, also inhibit DNA sensors cGAS and TLR9 demonstrating a broad utility against pathogenic DNA-driven inflammatory responses. The 4-sulfonic calixarenes inhibited AIM2 dependent post-stroke T cell death, highlighting a proof of concept that the 4-sulfonic calixarenes could be effective at combatting post-stroke immunosuppression. By extension, we propose a broad utility against DNA driven inflammation in disease. Finally, we reveal that the ancient drug suramin, by virtue of its structural similarities, is an excellent inhibitor of DNA-dependent inflammation and propose that suramin could be rapidly repurposed to meet an ever increasing clinical need.

## Introduction

Inflammation is part of the host response to protect against invading pathogens and to clear and repair tissue damage. Inflammation needs to be strictly regulated by the host as inappropriate inflammation leads to the development and worsening of disease(*1*). Inflammation is initiated when cells of the innate immune system detect damage associated molecular patterns (DAMPs) or pathogen associated molecular patterns (PAMPs) through activation of pattern recognition receptors (PRRs). Extranuclear double stranded (ds)DNA is an indicator of cellular damage or pathogen infection, and triggers a robust inflammatory response driven by the release of pro-inflammatory cytokines following the activation of absent in melanoma 2 (AIM2), cyclic GMP-AMP synthase (cGAS), and toll-like receptor 9 (TLR9)(*2*). Whilst dsDNA sensing has evolved to protect and alert the host to invading pathogens and damaged tissue, aberrant activation of nucleic acid-sensing pathways can lead to autoinflammatory disease. Mutations in DNA-sensing PRRs (e.g. cGAS) or regulators of DNA synthesis/degradation (e.g. DNase II), or excessive self-DNA release in the pathology of disease (e.g. myocardial infarction, stroke) can drive unwanted systemic inflammatory responses(*3*).

Inflammasomes are multi-protein complexes that drive inflammation in response to damage or infection. Inflammasome complexes are comprised of a soluble PRR associated with the adaptor protein ASC (apoptosis-associated speck-like protein containing a CARD), and the protease caspase 1. Inflammasome formation results in the processing and release of the pro-inflammatory cytokines IL-1β and IL-18, and the induction of lytic cell death by gasdermin D (GSDMD)-mediated pyroptosis(*4*). Inflammasome complexes are defined by their scaffolding PRR, with each one sensing particular PAMPs, DAMPs or cellular disturbances, including NLRP3, NLRC4 and the dsDNA sensor AIM2(*4*). Following detection of self, or non-self, cytosolic dsDNA, AIM2 forms an inflammasome(*5*). Whilst AIM2 has beneficial roles in promoting host defence responses, inappropriate activation of AIM2 is associated with worsening of diseases such as atherosclerosis(*6*), cancers such as melanoma(*7*), ischemic stroke(*8*, *9*), and post-stroke immunosuppression(*10*). We currently lack effective tools to pharmacologically block AIM2 responses, hindering investigation in pre-clinical research and clinical drug development.

Here, using *in vitro* cellular assays, *in silico* molecular modelling, and *in vivo* disease models, we have identified that 4-sulfonic calixarenes are potent inhibitors of dsDNA-driven inflammation through the AIM2 inflammasome by reversibly binding to the dsDNA-binding site. At higher doses, the 4-sulfonic calixarenes also inhibited the dsDNA sensors cGAS and TLR9. We demonstrate that 4-sulfonic calixarenes are effective *in vivo* at preventing AIM2-dependent post-stroke immunosuppression, providing the foundation for a new therapeutic approach to limit deadly infections post-stroke, as well as functioning as a new investigative tool to study dsDNA-dependent inflammatory processes in pre-clinical disease models. Finally, using the structure activity relationship identified with 4-sulfonic calixarenes, we used *in silico* approaches to identify clinically available analogues and established that the drug suramin is an effective inhibitor of the AIM2 inflammasome, highlighting future translational opportunities to target AIM2 in patients.

## Results

We set out to test the hypothesis that the poly-anionic nature of 4-sulfonic calix[6]arene could enable effective inhibition of the AIM2 inflammasome (Fig 1A). In a previous study, 4-sulfonic calix[6]arene was ineffective at inhibiting the NLRP3 inflammasome(*11*). To activate the AIM2 inflammasome we transfected LPS-primed murine bone marrow-derived macrophages (BMDMs) with the synthetic double stranded (ds)DNA sequence poly dA:dT(*12*) and measured cell death and IL-1β release. 4-Sulfonic calix[6]arene dose dependently inhibited AIM2-dependent cell death and IL-1β release (Fig 1B,C). 4-Sulfonic calix[6]arene had IC_50_ values of 2.20 μM and 3.00 μM for poly dA:dT-induced cell death and IL-1β release respectively (Fig 1B,C). Pre-incubation of 4-sulfonic calix[6]arene, but not the NLRP3 inflammasome inhibitor MCC950(*13*), also inhibited poly dA:dT-induced ASC oligomerisation and cleavage of caspase-1, IL-1β, and GSDMD in LPS-primed mouse BMDMs (Fig 1D). Furthermore, treatment with 4-sulfonic calix[6]arene did not inhibit IL-1β release in response to treatment with the NLRP3 activator silica(*14*), or transfection with flagellin to activate NLRC4(*15*) (Fig 1E,F), suggesting 4-sulfonic calix[6]arene does not inhibit NLRP3 or NLRC4. 4-Sulfonic calix[6]arene did not inhibit transfection efficacy per se as 4-sulfonic calix[6]arene had no effect on NLRC4 activation by flagellin transfection (Fig 1E,F, Supplementary Fig 1C), and had no effect on transfection of rhodamine-tagged poly dA:dT at inflammasome inhibiting concentrations (Supplementary Fig 1A, B).

**Figure 1.**
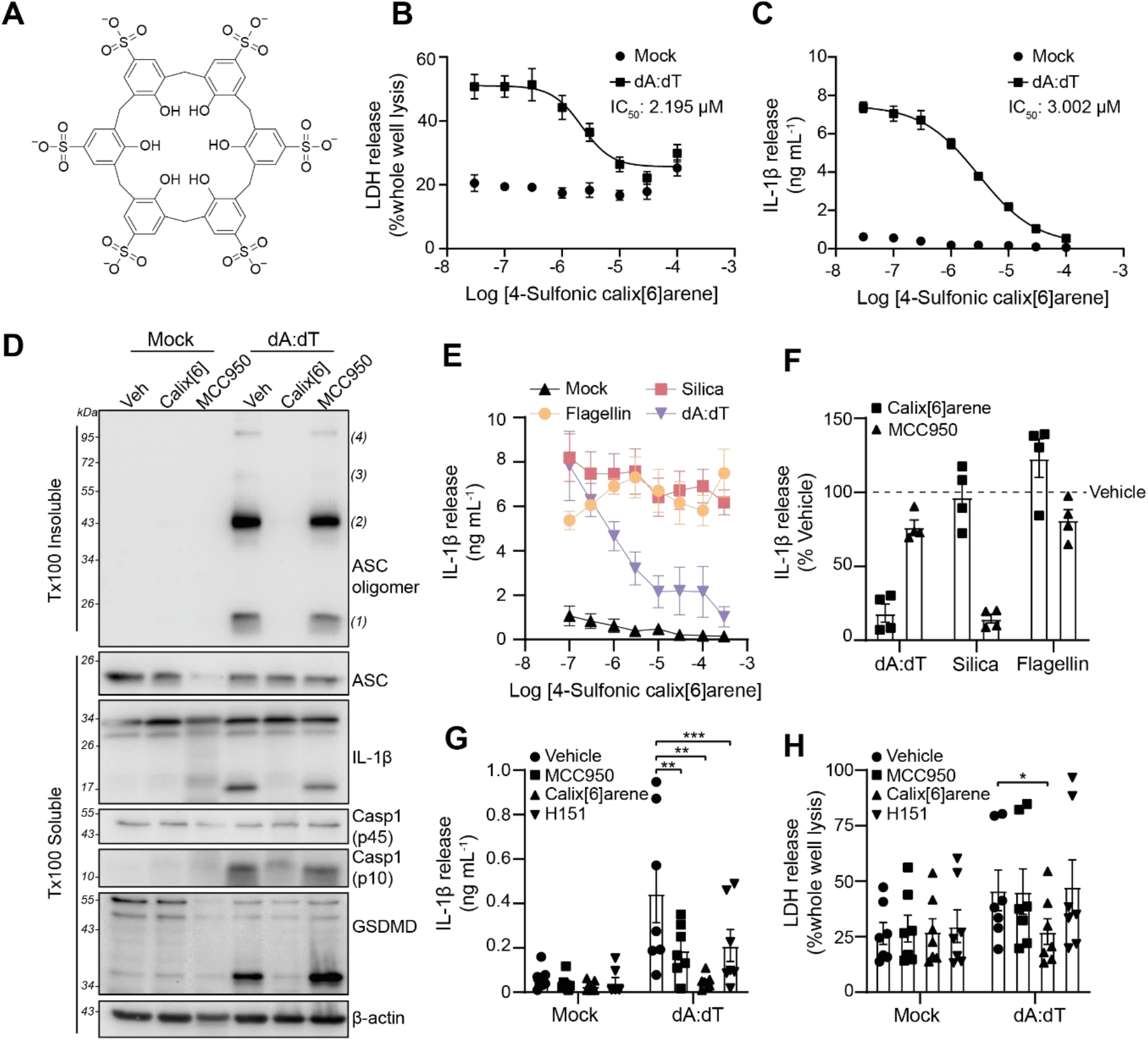
4-Sulfonic calix[6]arene inhibited double stranded (ds)DNA-induced inflammasome responses. **(A)** Chemical structure of 4-sulfonic calix[6]arene. **(B)** LDH and **(C)** IL-1β release in the supernatants of LPS-primed (1 μg mL^-1^, 4 h) bone marrow derived macrophages (BMDMs). BMDMs were pre-treated with the indicated concentration of 4-sulfonic calix[6]arene (0.03 - 100 μM) before transfection with poly dA:dT (1 μg mL^-1^, 4 h) (n=3). **(D)** Western blot of crosslinked ASC oligomers in Triton x100 (Tx100) insoluble BMDM cell lysates and corresponding ASC, IL-1β, Caspase-1 (Casp1) and GSDMD in the Tx100 soluble BMDM cell fraction. LPS-primed BMDMs were pre-treated with 4-sulfonic calix[6]arene (Calix[6], 30 μM), the NLRP3 inhibitor MCC950 (10 μM) or vehicle control (Veh, DMSO) before transfection with poly dA:dT (4 h) (n=4). **(E)** IL-1β release in the supernatant of LPS-primed BMDMs pre-treated with the indicated concentration of 4-sulfonic calix[6]arene (0.03 – 100 μM) before stimulation with silica (300 μg mL^-1^), flagellin (1 μg mL^-1^) or poly dA:dT (1 μg mL^-1^) for 4 h (n=4). **(F)** IL-1β release from LPS-primed BMDMs pre-treated with 4-sulfonic calix[6]arene (100 μM), MCC950 (10 μM) or vehicle control (DMSO) before stimulation with poly dA:dT (1 μg mL^-1^), silica (300 μg mL^-1^), or flagellin (1 μg mL^-1^) (n=4). **(G)** IL-1β and **(H)** LDH release in the supernatant from LPS-primed (1 μg mL^-1^, 4 h) human monocyte-derived macrophages (hMDMs) pre-treated with MCC950 (10 μM), 4-sulfonic calix[6]arene (30 μM), the STING inhibitor H151 (10 μM) or vehicle control (DMSO) before transfection with poly dA:dT (1 μg mL^-1^, 18 h) (n=7). Concentration-response curves were fitted using a four parameter logistical (4PL) model. *p<0.05, **p<0.01, ***p<0.001 determined by a two-way ANOVA with Dunnett’s post hoc analysis. Values shown are mean ± the SEM.

In human cells, the inflammasome response to dsDNA is reported to be dependent on a cGAS-STING-NLRP3 pathway rather than AIM2(*16*). In this model, it is proposed that dsDNA binds to and activates cGAS, resulting in synthesis of the cyclic messenger 2’,3’-cGAMP, which activates STING, leading to lysosomal rupture and activation of NLRP3(*16*). Therefore, we tested if 4-sulfonic calix[6]arene was effective at blocking dsDNA-induced inflammasome activation in LPS-primed human blood monocyte-derived macrophages (hMDMs). Matching previous reports(*16*), the NLRP3 inhibitor MCC950, and the STING inhibitor H151(*17*), partially inhibited IL-1β release from poly dA:dT transfected hMDMs (Fig 1G), and had no effect on cell death (Fig 1H). However, 4-sulfonic calix[6]arene completely inhibited dA:dT-transfection-induced IL-1β release and cell death (Fig 1G,H). These data indicate 4-sulfonic calix[6]arene inhibited dsDNA-induced inflammasome responses driven by both AIM2 and cGAS-STING.

We then examined the effect of 4-sulfonic calix[6]arene on the cGAS-STING pathway in more detail. Activation of cGAS-STING leads to a phosphorylation cascade including STING, TBK1 and IRF3 resulting in the transcription of type I interferons (IFNs)(*18*). 4-Sulfonic calix[6]arene caused a dose-dependent inhibition in poly dA:dT-induced IFN-β release from murine BMDMs, with an IC_50_ of 10.72 μM (Fig 2A). 4-Sulfonic calix[6]arene also reduced the phosphorylation of TBK1 and IRF-3 to a similar extent as the STING inhibitor H151 (Fig 2B-D). To test if 4-sulfonic calix[6]arene was acting via cGAS or STING, we used 10-carboxymethyl-9-acridanone (CMA), a small molecule that binds directly to, and activates murine STING independent of cGAS(*19*), and 2’,3’-cGAMP, the intracellular second messenger produced by cGAS that activates STING. CMA-induced IFN-β production in BMDMs was not significantly altered by pre-treatment with 4-sulfonic calix[6]arene, but was fully inhibited by H151 (Fig 2E). However, transfection of 2’,3’-cGAMP into BMDMs induced IFN-β that was inhibited by both 4-sulfonic calix[6]arene and H151 (Fig 2F), suggesting that 4-sulfonic calix[6]arene may be effective at both the dsDNA-binding site on cGAS and the 2’,3’-cGAMP, but not the CMA binding site on STING.

**Figure 2.**
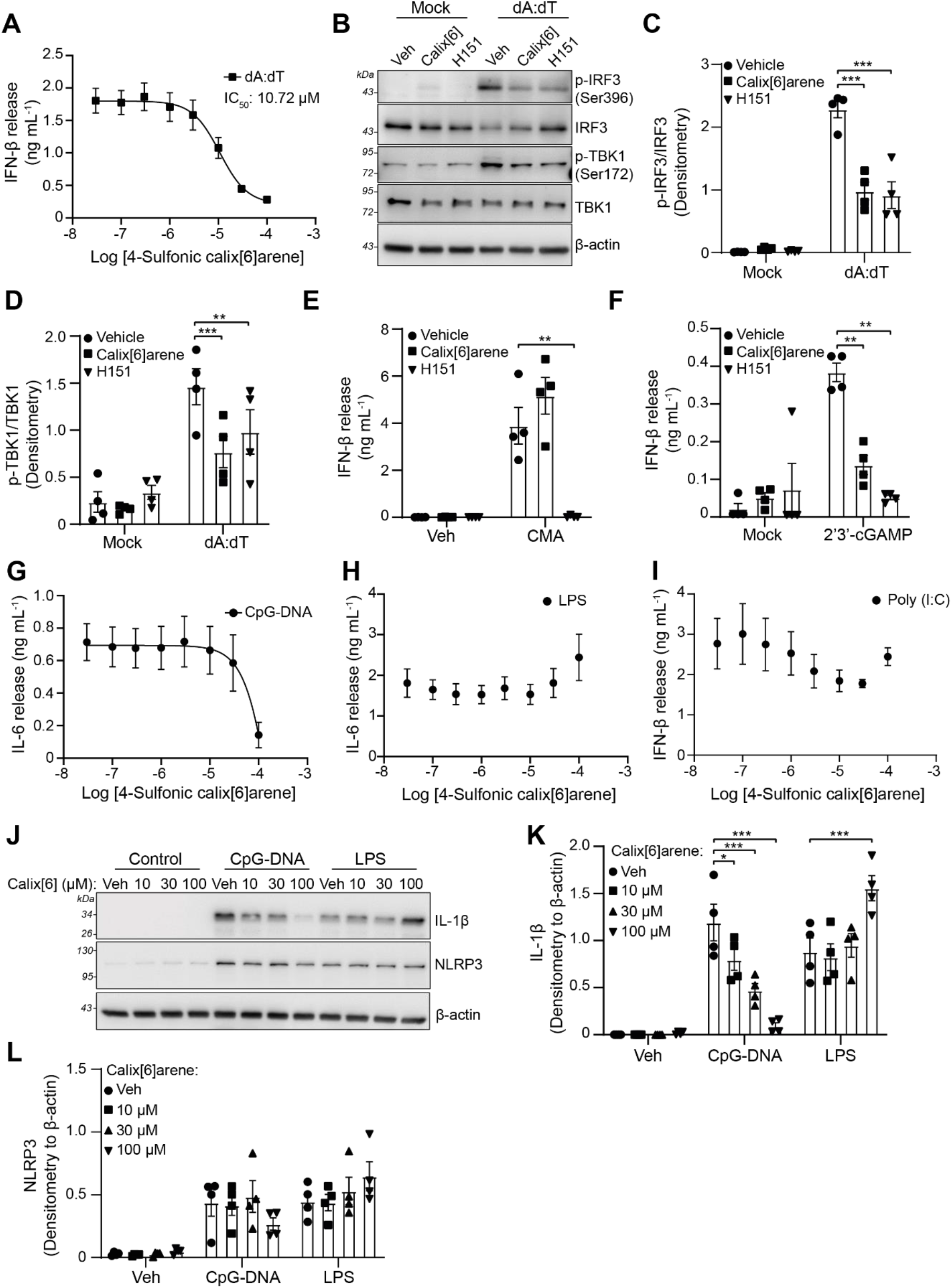
4-Sulfonic calix[6]arene inhibited dsDNA-induced type I interferon release and CpG DNA-induced inflammation. **(A)** IFN-β release in the supernatants of bone marrow derived macrophages (BMDMs). BMDMs were treated with the indicated concentration of 4-sulfonic calix[6]arene (0.03 – 100 μM) before transfection with poly dA:dT (1 μg mL^-1^, 6 h) (n=4). **(B)** Western blot of phosphorylated (p-) and total levels of IRF3 and TBK1 in BMDM lysates following treatment with 4-sulfonic calix[6]arene (calix[6], 30 μM) or the STING inhibitor H151 (10 μM) and transfected with poly dA:dT (1 μg mL^-1^, 3 h) (n=4). **(C)** The ratio of p-IRF3 (Ser396) to total IRF3 and **(D)** the ratio of p-TBK1 to total TBK1 determined by densitometry of experiments shown in (B) (n=4). **(E)** IFN-β release in the supernatants of BMDMs pre-treated with 4-sulfonic calix[6]arene (30 μM), H151 (10 μM) or vehicle control (DMSO) before stimulation with CMA (250 μg mL^-1^, 6 h) (n=4). **(F)** IFN-β release in the supernatants of BMDMs pre-treated with 4-sulfonic calix[6]arene (30 μM), H151 (10 μM) or vehicle control (DMSO) before transfection with 2’,3’-cGAMP (1.5 μg mL^-1^, 6 h) (n=4). **(G)** IL-6 release in the supernatants of BMDMs pre-treated with the indicated concentration of 4-sulfonic calix[6]arene (0.03 – 100 μM) and stimulated with CpG DNA (1 μM, 6 h) (n=4), **(H)** or LPS (1 μg mL^-1^, 6 h) (n=4). **(I)** IFN-β release in the supernatants of BMDMs pre-treated with the indicated concentration of 4-sulfonic calix[6]arene (0.03 – 100 μM) and stimulated with poly (I:C) (5 μg mL^-1^, 6h) (n=4). **(J)** Western blot of pro-IL-1β and NLRP3 in the lysates of BMDMs pre-treated with the indicated concentration of 4-sulfonic calix[6]arene, or vehicle control (Veh, DMSO) and stimulated with CpG DNA (1 μM, 6 h) or LPS (1 μg mL^-1^, 6 h) (n=4). **(K-L)** Densitometry of pro-IL-1β **(K)** and NLRP3 **(L)** from experiments in (J). Concentration-response curves were fitted using a four parameter logistical (4PL) model. *p<0.05, **p<0.01, ***p<0.001 determined by a two-way ANOVA with Dunnett’s post hoc analysis vs vehicle control. Values shown are mean ± the SEM.

We also examined if 4-sulfonic calix[6]arene was an inhibitor of the dsDNA sensing TLR9, an endosomal TLR that senses CpG-rich DNA(*20*). CpG-rich DNA is enriched in bacterial genomes and therefore TLR9 functions as a sensor for bacterial infection rather than host cell damage(*21*). Activation of TLR9 with CpG-rich DNA results in NF-κB signalling and pro-inflammatory gene expression, including IL-6, IL-1 and NLRP3(*22*, *23*). 4-Sulfonic calix[6]arene reduced CpG-induced IL-6 release in BMDMs at 100 μM, but was ineffective at lower concentrations (Fig 2G). 4-Sulfonic calix[6]arene was not effective up to 100 μM on LPS-induced TLR4, or poly (I:C)-induced TLR3 responses (Fig 2H,I). 4-Sulfonic calix[6]arene pre-treatment also dose-dependently inhibited CpG-induced pro-IL-1β expression, but not in response to LPS (Fig 2J,K). LPS- or CpG-induced expression of NLRP3 was not prevented by pre-treatment with 4-sulfonic calix[6]arene (Fig 2J,L). These data suggest that 4-sulfonic calix[6]arene can inhibit TLR9 receptors, but at much higher concentrations than is required for inhibition of AIM2 and cGAS.

That 4-sulfonic calix[6]arene was effective at inhibiting a range of dsDNA sensors suggested that a potential mechanism of action could be through interacting with the dsDNA binding sites. 4-Sulfonic calix[6]arene is negatively charged, resulting from the six sulfonic acid groups, a property that is similar to the phosphate backbone of dsDNA. The negative charges of the DNA backbone are essential for its binding to DNA sensors such as AIM2(*24*) and cGAS(*25*). To investigate if the negatively charged sulfonic acid groups contributed to the inhibitory actions of 4-sulfonic calix[6]arene on dsDNA sensors, we tested the calixarene compounds: 4-sulfonic calix[4]arene, 4-sulfonic calix[8]arene and 4-*tert-*butyl calix[6]arene (Fig 3A). 4-Sulfonic calix[4]arene and 4-sulfonic calix[8]arene contain two fewer and two extra sulfonic acid groups than 4-sulfonic calix[6]arene respectively, and exhibit corresponding differences in charge. 4-*tert*-Butyl calix[6]arene exhibits the same number of phenolic units as 4-sulfonic calix[6]arene but contains neutral *tert*-butyl groups on each phenolic unit instead of negative sulfonic acid groups. Comparisons between 4-sulfonic calix[6]arene and 4-*tert*-butyl calix[6]arene would therefore determine if the inhibitory action of 4-sulfonic calix[6]arene required the sulfonic acid groups and negative charge. LPS-primed BMDMs were treated with increasing concentrations of either 4-sulfonic calix[4]arene, 4-sulfonic calix[6]arene, 4-sulfonic calix[8]arene or 4-*tert*-butyl calix[6]arene before transfection with poly dA:dT to activate AIM2. 4-*tert*-Butyl calix[6]arene was unable to prevent poly dA:dT-induced cell death or IL-1β release when used up to a concentration of 100 μM (Fig 3B,C). All 4-sulfonic calixarenes inhibited AIM2-dependent IL-1β release, and the potency of inhibition correlated with the number of sulfonate groups, with 4-sulfonic calix[4]arene the least effective and 4-sulfonic calix[8]arene the most effective (Fig 3B). 4-Sulfonic calix[8]arene was highly potent at inhibiting IL-1β release (IC_50_:214 nM) (Fig 3B), cell death (IC_50_:191 nM) (Fig 3C), ASC oligomerisation, and cleavage of IL-1β, caspase-1 and GSDMD (Fig 3D). Pre-treatment with 4-sulfonic calix[4]arene or 4-*tert*-butyl calix[6]arene did not inhibit the formation of AIM2 inflammasomes up to 100 μM (Fig 3D). We then sought to identify if 4-sulfonic calixarenes were reversible inhibitors of the AIM2 inflammasome. LPS-primed BMDMs were incubated with a vehicle or 30 μM of each 4-sulfonic calixarene for 1 hour and then washed three times before transfection with poly dA:dT and assessment of AIM2 activation by IL-1β release. Unwashed BMDMs and BMDMs treated with additional 4-sulfonic calixarenes after the washes were used in parallel as controls. Consistent with the concentration-response data, 4-sulfonic calix[6]arene and 4-sulfonic calix[8]arene were effective at inhibiting AIM2-dependent IL-1β release in unwashed BMDMs, whereas 4-sulfonic calix[4]arene was inactive (Fig 3E). The inhibition of AIM2 by 4-sulfonic calix[6]arene and 4-sulfonic calix[8]arene was completely lost following washing, and was restored upon re-addition of the respective compound (Fig 3E). These data indicate that inhibition of AIM2 by 4-sulfonic calixarenes was reversible.

**Figure 3.**
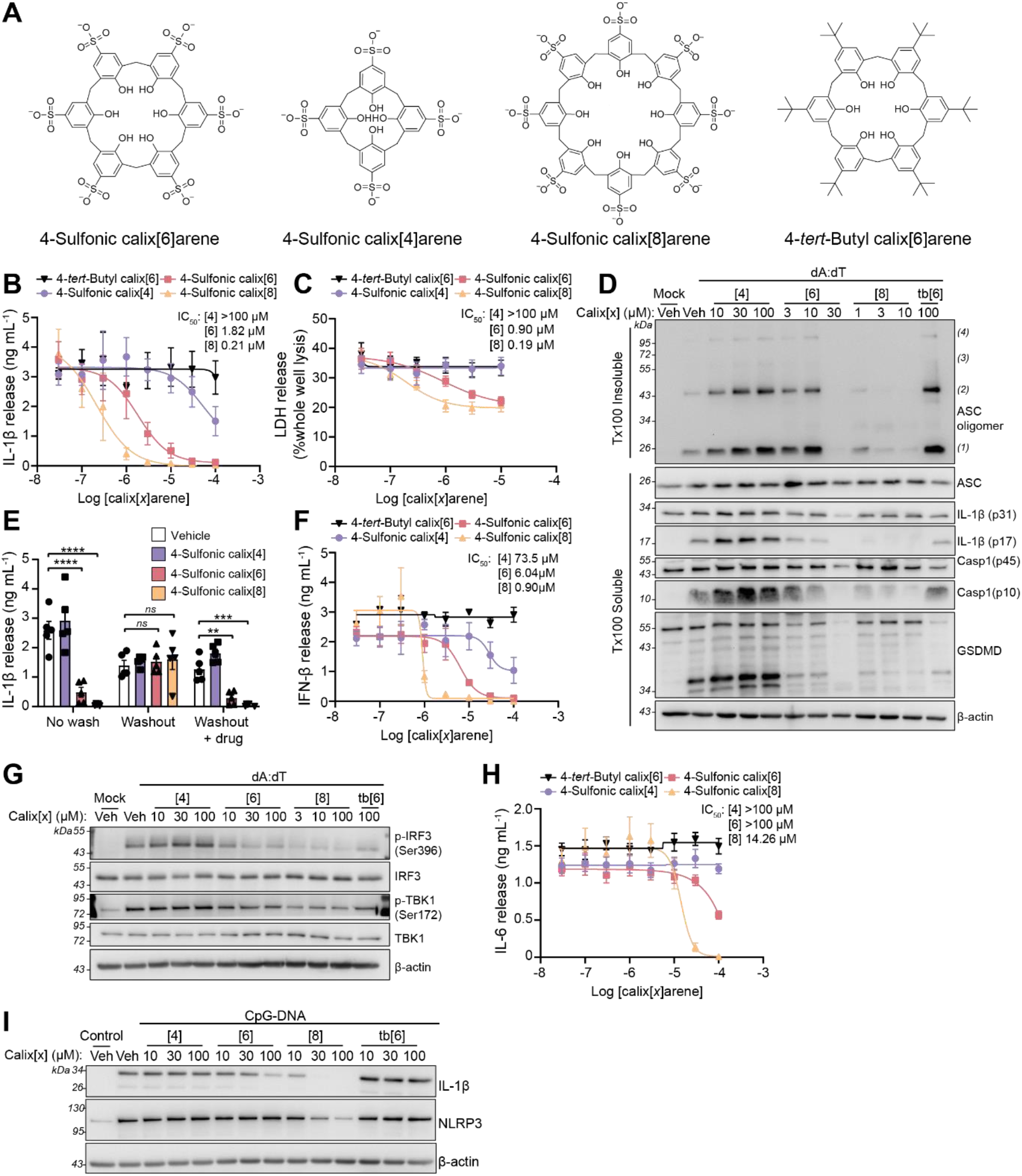
Inhibition of dsDNA inflammatory signalling by 4-sulfonic calixarenes is readily reversible and is dependent on the sulfonic acid groups. **(A)** Chemical structure of 4-sulfonic calix[6]arene, 4-sulfonic calix[4]arene, 4-sulfonic calix[8]arene and 4-*tert*-butyl calix[6]arene. **(B)** IL-1β release and **(C)** LDH release in the supernatants of LPS-primed (1 μg mL^-1^, 4 h) bone marrow derived macrophages (BMDMs). BMDMs were pre-treated with the indicated concentration of 4-sulfonic calix[4]arene, 4-sulfonic calix[6]arene, 4-sulfonic calix[8]arene or 4-*tert*-butyl calix[6]arene (0.03 - 100 μM) before transfection with poly dA:dT (1 μg mL^-1^, 4 h) (n=4). **(D)** Western blot of crosslinked ASC oligomers in Triton x100 (Tx100) insoluble BMDM cell lysates and corresponding ASC, IL-1β, Caspase-1 (Casp1) and GSDMD in the Tx100 soluble BMDM cell fraction. LPS-primed BMDMs were pre-treated with indicated concentrations of 4-sulfonic calix[4]arene ([4]), 4-sulfonic calix[6]arene ([6]), 4-sulfonic calix[8]arene ([8]), 4-*tert*-butyl calix[6]arene (tb[6]), or vehicle control (Veh, DMSO) before transfection with poly dA:dT (4 h) (n=4). **(E)** IL-1β release in to the supernatants of LPS-primed (1 μg mL^-1^) BMDMs, treated with 30 μM 4-sulfonic calix[4]arene, 4-sulfonic calix[6]arene, 4-sulfonic calix[8]arene, or vehicle control (DMSO) before transfection with poly dA:dT (1 μg mL^-1^, 3.5 h) (n=5). BMDMs were either stimulated in 4-sulfonic calixarene containing media (no wash), washed 3x before stimulation (washout), or washed 3x with re-addition of each respective drug (washout + drug). **(F)** IFN-β release into the supernatants of BMDMs. BMDMs were treated with the indicated concentration of 4-sulfonic calix[4]arene, 4-sulfonic calix[6]arene, 4-sulfonic calix[8]arene or 4-*tert*-butyl calix[6]arene (0.03 – 100 μM) before transfection with poly dA:dT (1 μg mL^-1^, 6 h) (n=4). **(G)** Western blot of phosphorylated (p-) and total levels of IRF3 and TBK1 in BMDM lysates following treatment with indicated concentrations of 4-sulfonic calix[4]arene ([4]), 4-sulfonic calix[6]arene ([6]), 4-sulfonic calix[8]arene ([8]), 4-*tert*-butyl calix[6]arene (tb[6]), or vehicle control (Veh, DMSO) before transfection with poly dA:dT (3 h) (n=4). **(H)** IL-6 release in the supernatants of BMDMs pre-treated with the indicated concentration of 4-sulfonic calix[4]arene, 4-sulfonic calix[6]arene, 4-sulfonic calix[8]arene or 4-*tert*-buyl calix[6]arene (0.03 – 100 μM) and stimulated with CpG DNA (1 μM, 6 h) (n=4). **(I)** Western blot of pro-IL-1β and NLRP3 in the lysates of BMDMs pre-treated with indicated concentrations of 4-sulfonic calix[4]arene ([4]), 4-sulfonic calix[6]arene ([6]), 4-sulfonic calix[8]arene ([8]), 4-*tert*-butyl calix[6]arene (tb[6]), or vehicle control (Veh, DMSO) and stimulated with CpG DNA (1 μM, 6 h) (n=4). Concentration-response curves were fitted using a four parameter logistical (4PL) model. *p<0.05, **p<0.01, ***p<0.001 determined by a two-way ANOVA with Dunnett’s post hoc analysis vs vehicle control. Values shown are mean ± the SEM.

We next tested the additional calixarene compounds on preventing activation of cGAS-STING and TLR9. Similar to AIM2 inhibition, 4-*tert*-butyl calix[6]arene was unable to prevent cGAS-STING-dependent IFN-β release and 4-sulfonic calixarenes inhibited at a potency correlated to the number of sulfonates (Fig 3F). 4-Sulfonic calix[8]arene was the most potent (IC_50_: 901 nM) (Fig 3F) and also inhibited cGAS-STING-dependent phosphorylation of TBK-1 and IRF-3 at 3 μM (Fig 3G, Supplementary Fig 2A,B). Similarly, CpG-induced TLR9 responses were not inhibited by 4-*tert*-butyl calix[6]arene or 4-sulfonic calix[4]arene, but were inhibited by 4-sulfonic calix[6]arene and 4-sulfonic calix[8]arene. 4-Sulfonic calix[8]arene was the most potent, inhibiting CpG-induced IL-6 release (IC_50_: 14.12 μM) (Fig 3H), and induction of IL-1β and NLRP3 expression (Fig 3I, Supplementary Fig 2C,D). These data suggest that reversible inhibition of AIM2, and inhibition of cGAS and TLR9, by 4-sulfonic calixarenes is driven by the negatively charged sulfonic acid groups.

Next, molecular modelling was used to assess if 4-sulfonic calixarenes would interact with the dsDNA binding site of the HIN domain of AIM2. The X-ray co-crystal structure of human AIM2 bound to dsDNA (PDB:3RN5, (*24*) resolution 2.50 Å) showed 4 identical HIN domains with bound dsDNA (Fig 4A, Supplementary Fig 3). To validate the model, dsDNA was removed and the dsDNA phosphate backbone containing only the ribose sugars (i.e. with bases replaced with H) was docked into the AIM2 HIN domain between the oligonucleotide binding sites OB1 and OB2 using Autodock Vina(*26*). This showed a good overlay between the docked and co-crystallised dsDNA (Fig 4A). The electrostatics of the binding of docked dsDNA in AIM2 HIN shows that the negatively charged phosphate groups are appropriately placed with respect to the positively charged lysine and arginine side chains of the HIN domain (Fig 4B,C).

**Figure 4.**
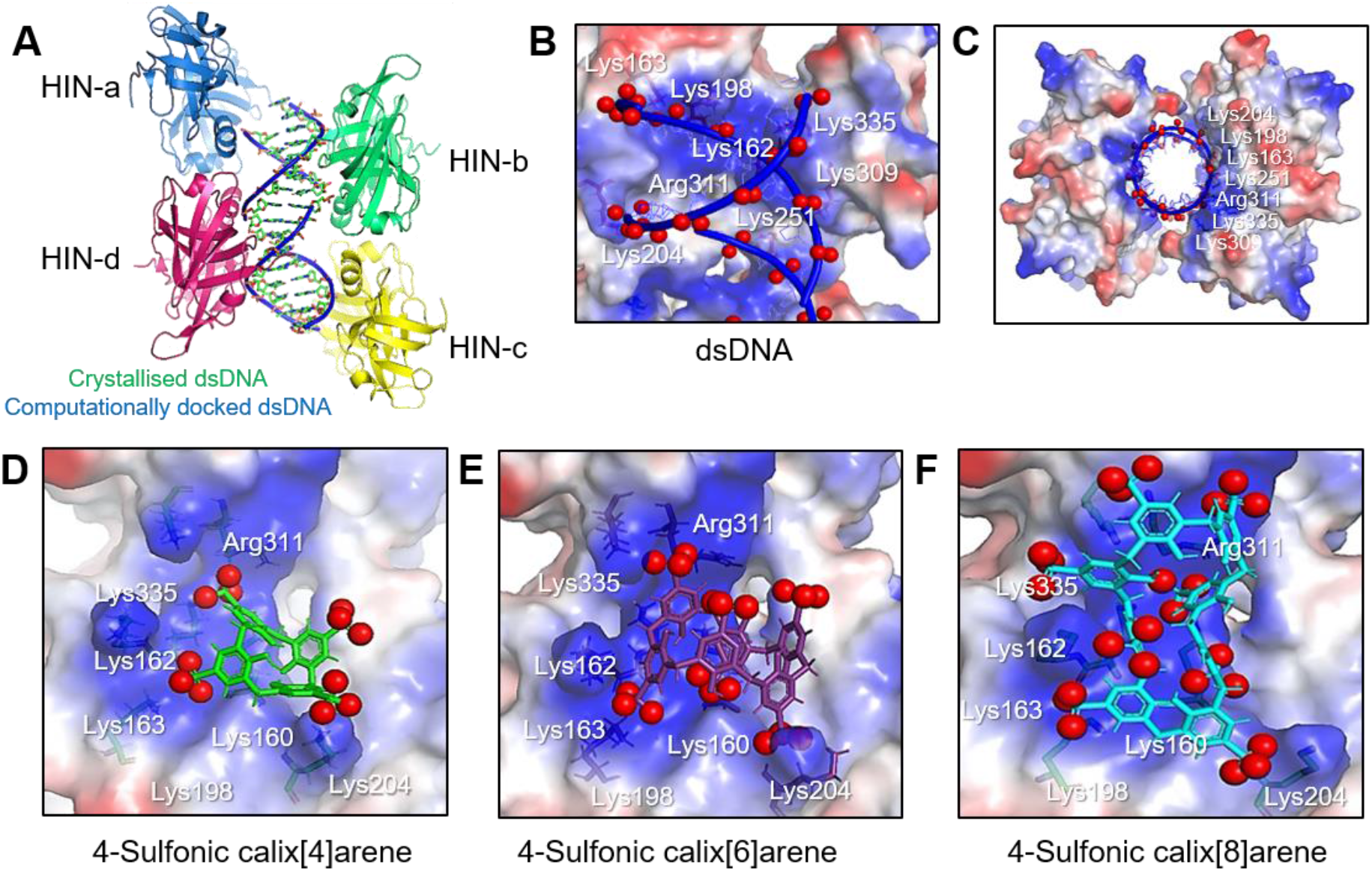
Modelling of 4-sulfonic calixarenes on the X-ray crystal structure of dsDNA bound AIM2 HIN domains. **(A-C)** Docking validation in human AIM2 HIN domain (PDB:3RN5), **(A)** redocked dsDNA backbone and ribose sugar (blue) overlaid on co-crystallized dsDNA (green) showing the major groove regions (red dashes) and the minor groove regions (brown dashes) of dsDNA, **(B-C)** electrostatic potential maps for docked dsDNA showing interacting amino acid residues with negatively charged phosphate groups (red spheres), **(B)** side view **(C)** top view. **(D-F)** Electrostatic potential maps for docked 4-sulfonic calix[*x*]arenes in the human AIM2 HIN domain showing interacting amino acid side chains with negatively charged sulfonate oxygen atoms (red spheres) for **(D)** 4-sulfonic calix[4]arene, **(E)** 4-sulfonic calix[6]arene and **(F)** 4-sulfonic calix[8]arene. For electrostatic potential maps: blue: positive region, red: negative region, white: neutral region, red spheres: negatively charged atoms for the phosphates of dsDNA (A-C) or sulfonate oxygen atoms (D-E). Images created using Pymol.

Previous reports have shown that 4-sulfonic calixarenes are flexible with 4-sulfonic calix[4]arene, 4-sulfonic calix[6]arene and 4-sulfonic calix[8]arene adopting 4, 8 and 16 conformations, respectively(*27*–*31*). A conformational search was carried out for 4-sulfonic calixarenes using Molecular Operating Environment (MOE 2015.08, Chemical Computing Group, Canada) (Supplementary Table 1, Supplementary Fig 4), with the lowest energy conformer for 4-sulfonic calix[4]arene adopting the partial cone, for 4-sulfonic calix[6]arene the 1,2,4-alternate and for 4-sulfonic calix[8]arene the 1,2,4,6,8-alternate. The top 49 conformers of each 4-sulfonic calixarene (Supplementary Table 1) were docked flexibly in the human AIM2 HIN domain (monomer) (PDB:3RN5)(*24*) using MOE and the lowest energy conformers were docked rigidly using Autodock Vina(*26*). The predominant binding mode of 4-sulfonic calix[4]arene in the AIM2 HIN domain (monomer) is the 1,2-alternate conformation (docking score −5.0 Kcal/mol) in MOE, which showed the greatest number of interactions of the sulfonate groups with the HIN domain residues (Fig 4D, Supplementary Tables 1, 2). This finding is consistent with docking using Autodock Vina (Supplementary Table 1, Table 1). Low energy docking scores were also found for the binding of 4-sulfonic calix[6]arene to AIM2 using MOE, with the best conformation being the 1,3-alternate (docking score: −8.4 Kcal/mol) showing docking interactions similar to those observed with the DNA phosphate groups (Fig 4E, Supplementary Tables 1 & 2). Docking with Autodock Vina also gave comparable results (Table 1, Supplementary Table 3). Consistent with its lower experimental IC_50_ value (and thus greater potency), 4-sulfonic calix[8]arene docking showed strong binding to AIM2 forming six ionic interactions, compared to 3 and 5 for the calix[4] and calix[6] analogues, respectively (Supplementary Table 1). The preferred docked conformation of 4-sulfonic calix[8]arene using MOE was the 1,2,4,6,8-alternate with a docking score of −9.9 Kcal/mol (Fig 4F) (Supplementary Table 2). Similar results were observed using Autodock Vina (Supplementary Table 3). The docked structure shows that six of the eight sulfonate groups were involved in intermolecular interactions (Table 1). Another possible explanation why 4-sulfonic calix[8]arene is the most active is the largest measured distance between the two sulfonate oxygen atoms of 21.4 Å, similar to the distance between 2 oxygen atoms of co-crystallised dsDNA in the minor groove (20.4 Å) (Supplementary Fig 5A)(*24*). This supports 4-sulfonic calix[8]arene competing with DNA in the HIN domain interface (Supplementary Fig 5B). Equivalent distances are much shorter for 4-sulfonic calix[4]arene and 4-sulfonic calix[6]arene, being 13.5 Å and 15.9 Å, respectively (Supplementary Fig 5C-D). Summarising this molecular modelling, the 4-sulfonic calixarenes showed excellent binding in the AIM2 HIN domain with the sulfonic acid groups binding in a similar way to the phosphate groups of dsDNA, confirming that they are convincing dsDNA mimetics. The larger 4-sulfonic calix[8]arene showed the best binding score, with six of the eight sulfonic acid groups binding to AIM2.

**Table 1:** Distances for the ionic interactions and docking scores of the best conformers for 4-sulfonic calixarenes in the human AIM2 HIN domain (monomer) (PDB:3RN5) using MOE & Autodock Vina. Distances are from the sulfonate oxygen to the positively charged nitrogen atoms in the HIN domain binding pocket, or from the negatively charge phosphate oxygen in co-crystallized dsDNA. *Sulfonate oxygen atoms form a hydrogen bond with the amino acid NH/CH backbone.

| Interacting residue | Co-crystallized dsDNA | MOE (Å) |  |  | Autodock Vina (Å) |  |  |  |
| --- | --- | --- | --- | --- | --- | --- | --- | --- |
|  |  | 4-Sulfonic calix[4]arene | 4-Sulfonic calix[6]arene | 4-Sulfonic calix[8]arene | Redocked dsDNA | 4-Sulfonic calix[4]arene | 4-Sulfonic calix[6]arene | 4-Sulfonic calix[8]arene |
| Docking score (Kcal/mol) | - | -5.0 | -8.4 | -9.9 | - | -11.7 | -12.4 | -13.3 |
| Lys160 | - | 3.0 | 3.1 (OH) | 3.2 | - | 3.2 | 2.9 (OH) | 3.0 |
| Lys162 | 3.4 | - | - | 3.2 | 3.0 | - | - | 3.8 |
| Lys163 | 3.0 | - | 3.0 | 3.0 | 3.0 | - | 3.0 | 3.5 |
| Lys198 | 3.3 | - | 3.4 | 3.3 | 3.0 | - | 3.4 | 3.1 |
| Lys204 | 3.3 | 3.0 | 2.9 | 3.0 | 3.0 | 3.1 | 3.2 | 3.7 |
| Arg311 | 2.8 | 3.7 | 3.1 | 3.3 | 4.0 | 3.1 | 3.3 | 3.8 |
| Lys335 | 3.0 | - | 2.9 | 3.0* | 3.8 | - | 3.1 | 3.3* |

Dysregulation of extracellular and cytosolic dsDNA is implicated in the pathophysiology of various diseases, including autoimmune disease, cancer, and stroke(*32*, *33*). We therefore hypothesised that 4-sulfonic calixarenes could be used as therapeutic or research tools to investigate the consequences of dsDNA signalling in disease. The AIM2 inflammasome mediates post-stroke immunosuppression which predisposes individuals to systemic infections(*10*). Infections are a major cause of death post-stroke(*34*), highlighting the need to find therapeutic approaches to limit post-stroke immunosuppression. Brain injury results in an increase in dsDNA in the blood, causing AIM2 inflammasome activation in blood monocytes and IL-1β release, which subsequently induces FAS ligand (FASL)-FAS-dependent T cell apoptosis leading to immunosuppression(*10*). We therefore tested if 4-sulfonic calix[6]arene prevented AIM2-dependent induction of T-cell death following stroke. Whole splenocytes were isolated from naïve mice and treated with 4-sulfonic calix[6]arene before incubation with serum (25% v/v) from sham- or stroke-operated mice for 16 hours. Treatment of splenocytes with serum from stroke-operated mice, which contains increased levels of dsDNA, resulted in a loss of T cells determined by flow cytometry, and this was prevented by 4-sulfonic calix[6]arene (Fig 5A,B). We also used a co-culture of BMDMs and T cells, to test if stroke serum-induced T cell death was blocked by co-treatment with 4-sulfonic calix[6]arene. LPS-primed BMDMs were incubated with serum (25% v/v) from sham- or stroke-operated mice for 10 minutes, before the serum was washed off and murine T cells were added to the culture for 4 h. In agreement with the whole splenocyte culture, T cell death was increased following incubation with serum from stroke-operated mice which was inhibited by pre-treatment with 4-sulfonic calix[6]arene for 1 h (Fig 5C,D). To further examine the potential of 4-sulfonic calix[6]arene as a pharmacological tool to inhibit AIM2-dependent responses *in vivo*, we assessed T cell loss after stroke in vehicle or 4-sulfonic calix[6]arene-treated mice (Fig 5E). The splenic T cell count was significantly increased in 4-sulfonic calix[6]arene treated animals post-stroke compared to untreated (Fig 5F). Furthermore, in spleens of animals treated with 4-sulfonic calix[6]arene there was less caspase-1 activation post-stroke, determined by western blotting for cleaved caspase-1 (Fig 5G,H) and by FLICA (Fig 5I), suggesting an inhibition of inflammasome activation. These results confirm that 4-sulfonic calix[6]arene is also efficient *in vivo* blocking stroke-induced AIM2 inflammasome activation and thereby preventing post-stroke T cell death. This therapeutic use of 4-sulfonic calix[6]arene to prevent injury-induced immunosuppression via AIM2 activation could also be generalizable to other diseases of sterile tissue injury for which similar mechanisms have been demonstrated such as skin burn injury(*10*).

**Figure 5.**
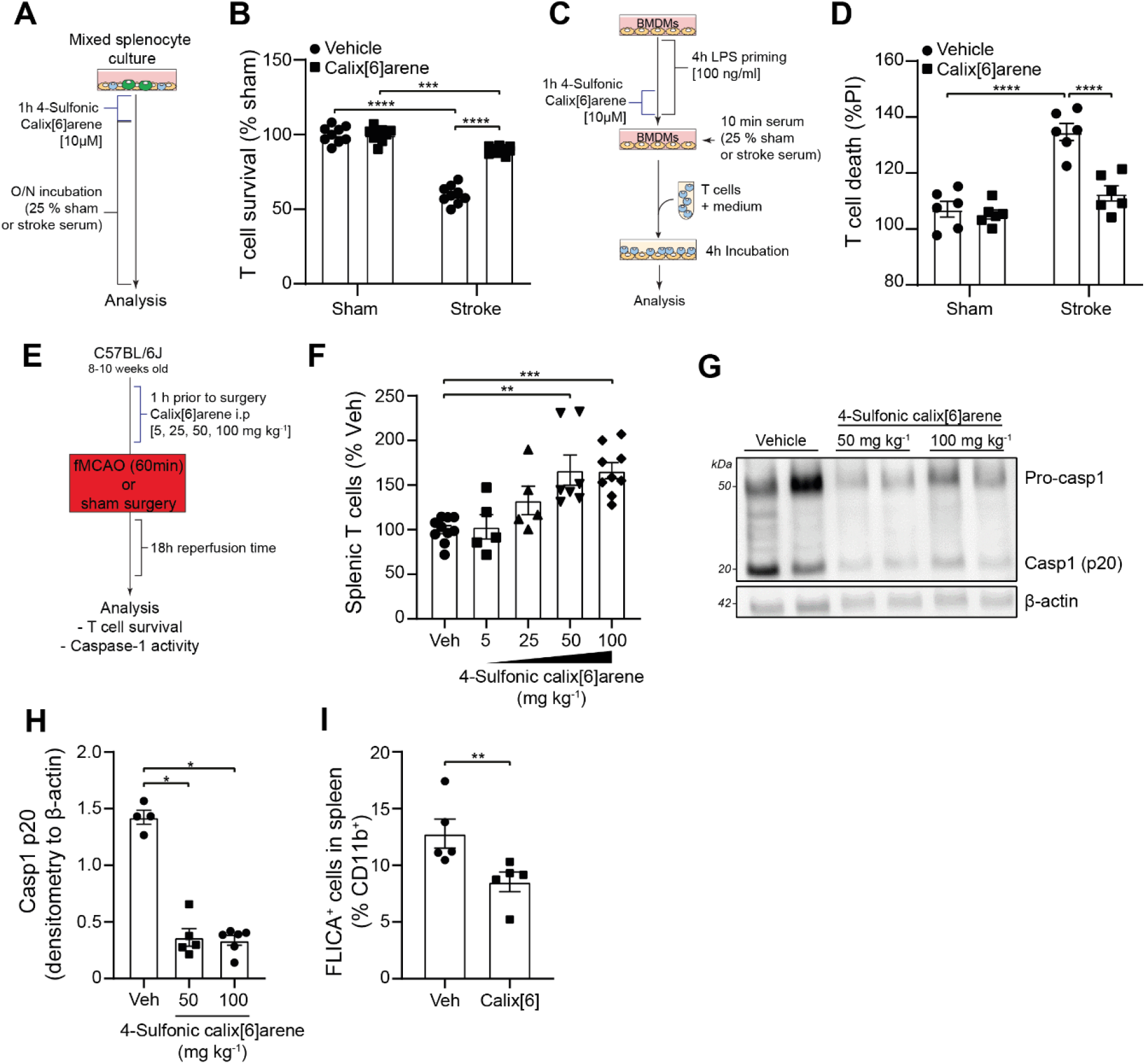
4-Sulfonic calix[6]arene inhibits AIM2 inflammasome-dependent T-cell death in a murine model of experimental stroke. **(A)** Schematic of the mixed splenocyte culture experiment: Whole splenocytes were pretreated with 4-sulfonic calix[6]arene (10 μM, 1 h) and then stimulated with serum (25 % v/v) from sham- or stroke-operated mice for 16 h. **(B)** CD3^+^ T cell survival in mixed splenocyte culture assessed by flow cytometry, shown as survival of T cells normalised to the sham serum-treated group (n=9, 3 independent experiments). **(C)** Schematic for the BMDM-T cell co culture experiment: BMDMs were LPS primed (100 ng ml^-1^, 4 h), treated with 4-sulfonic calix[6]arene (10 μM, 1 h) and stimulated with 25 % (v/v) serum of sham- or stroke-operated mice. Afterwards T cells were added for 4 h before T cell death was assessed by propidium iodide (PI) uptake. **(D)** T cell death assessment by flow cytometry from BMDM-T cell co cultures. T cell death is shown as percentage of PI uptake normalised to untreated co culture (n=6, 3 independent experiments). **(E)** Schematic of murine filamentous middle cerebral artery occlusion (fMCAO) experiment: mice received a intraperitoneal (i.p) injection of 5, 25, 50 or 100 mg kg^-1^ 4-sulfonic calix[6]arene 1 h prior to a stroke. After 18 h of reperfusion time the mice were sacrificed and analysed. **(F)** T cell survival after experimental ischemic stroke. Splenic T cell counts were analysed by flow cytometry, here shown as percentage of T cells in the vehicle group (n=5-9). **(G)** Representative caspase-1 western blot (Pro and p20 subunit) of whole splenocyte lysates from post-stroke mice in (F). **(H)** Quantification of caspase-1 p20 subunit normalized to β-actin from whole splenocyte lysates of vehicle (Veh, saline) and 4-sulfonic calix[6]arene (50 or 100 mg kg^-1^)–treated mice (n=4-6). **(I)** Splenic caspase-1 activity post-stroke measured by FAM660 FLICA. Mice were administered 4-sulfonic calix[6]arene (100 mg kg-1, i.p.), or vehicle (Veh, saline), 1 hour prior to fMCAO (n=5). *p<0.05, **p<0.01, ***p<0.001, ****p<0.0001 determined by a Kruskal-Wallis test with Dunnett’s post hoc analysis (B,D,F,H), or a Mann-Whitney U test (I). Values shown are mean ± the SEM.

Inhibition of dsDNA sensors by 4-sulfonic calixarenes is mediated by the multiple negatively charged sulfonic acid groups, leading us to examine whether any clinically available drugs exhibited similar properties and have the potential to be effective inhibitors of dsDNA-induced inflammation. The polyanionic drug suramin(*35*), used since the 1920s to treat African sleeping sickness and river blindness(*35*), also described as a purinergic receptor antagonists(*36*, *37*), was identified as a potential compound with similar properties to 4-sulfonic calixarenes due to the multiple sulfonic acid groups (Fig 6A). Suramin is already proposed to inhibit several dsDNA-binding proteins, including cGAS(*38*), Mcm10(*39*) and DNA topoisomerase II(*40*), but it is unknown whether suramin could inhibit the AIM2 inflammasome. Suramin pre-treatment of LPS-primed BMDMs dose-dependently inhibited AIM2 inflammasome activation, with an IC_50_ of 1.45 μM and 1.48 μM on AIM2-dependent LDH and IL-1β release respectively (Fig 6B,C). Furthermore, suramin inhibited AIM2-dependent ASC oligomerisation, caspase-1 and IL-1β processing (Fig 6D). Suramin did not inhibit ASC-oligomerisation, caspase-1 or IL-1β processing in response to stimulation of the NLRP3 or NLRC4 inflammasome with silica or flagellin respectively (Fig 6D). The inhibitory effect of suramin on AIM2 was reversible, as IL-1β release could be fully restored by repeated washing of suramin-treated BMDMs (Fig 6E). Similar to 4-sulfonic calix[6]arene and 4-sulfonic calix[8]arene, suramin was also effective at inhibiting cGAS-STING and TLR9 mediated inflammatory responses (Supplementary Fig 6A,B). Suramin and 4-sulfonic calix[8]arene also inhibited the dsDNA-induced inflammasome response in human macrophages blocking both pyroptosis (Fig 6F) and IL-1β release (Fig 6G). These data identify suramin as a clinically available drug that reversibly inhibits the AIM2 inflammasome in human and mice.

**Figure 6.**
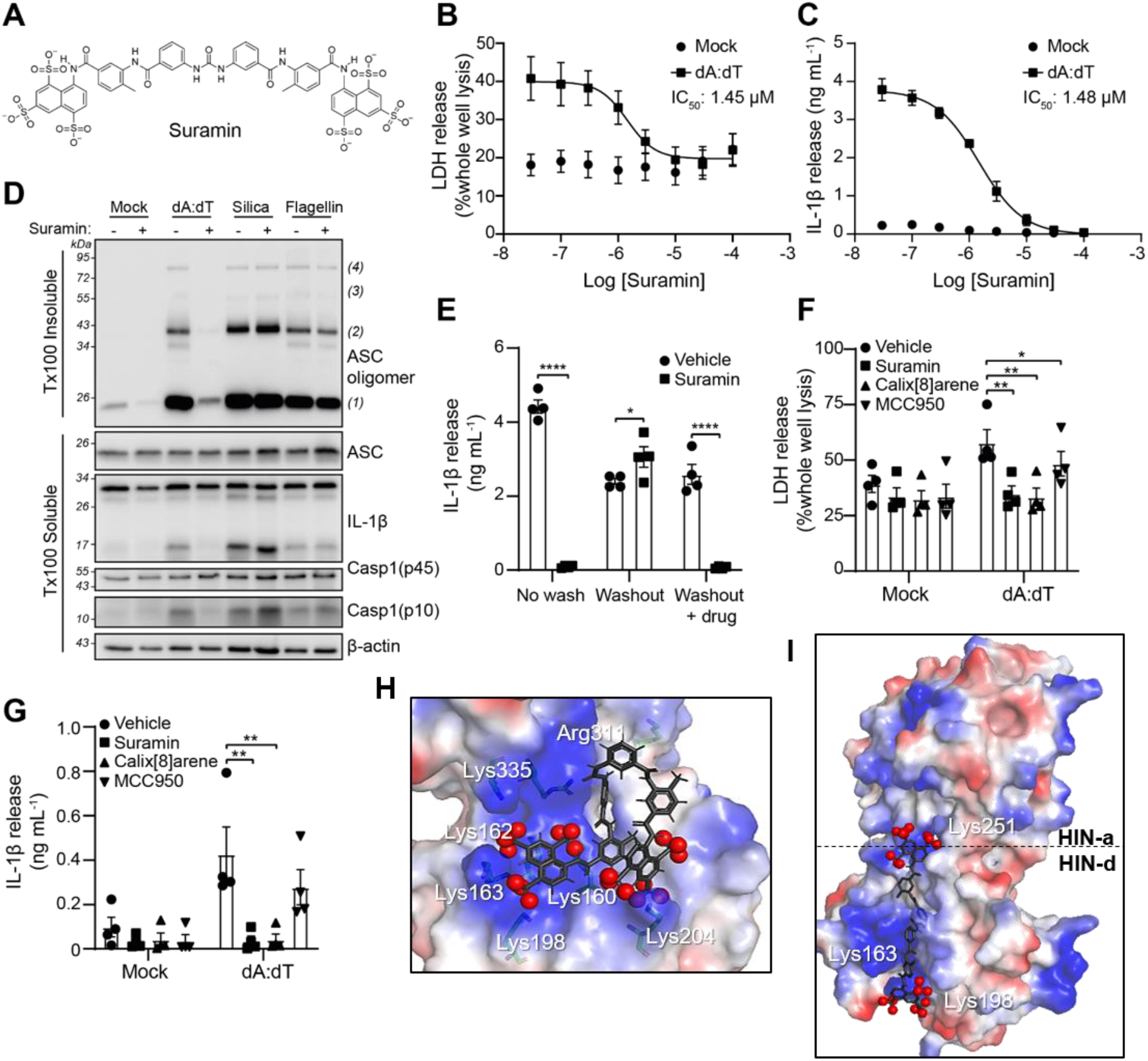
Suramin is an inhibitor of the AIM2 inflammasome. **(A)** Chemical structure for suramin. **(B)** LDH and **(C)** IL-1β release in the supernatants of LPS-primed (1 μg mL^-1^, 4 h) bone marrow derived macrophages (BMDMs). BMDMs were pre-treated with the indicated concentration of suramin (0.03 - 100 μM) before transfection with poly dA:dT (1 μg mL^-1^, 4 h) (n=4). **(D)** Western blot of crosslinked ASC oligmers in Triton x100 (Tx100) insoluble BMDM cell lysates and corresponding ASC, IL-1β, Caspase-1 (Casp1) and GSDMD in the Tx100 soluble BMDM cell fraction. LPS-primed BMDMs were pre-treated with Suramin (10 μM) or vehicle control (Veh, DMSO) before transfection with poly dA:dT (1 μg mL^-1^), transfection of flagellin (1 μg mL^-1^), or treatment with silica (300 μg mL^-1^) for 4 h (n=4). **(E)** IL-1β release in the supernatants of LPS-primed (1 μg mL^-1^) BMDMs, treated with suramin (10 μM), or vehicle control (DMSO) before transfection with poly dA:dT (1 μg mL^-1^, 3.5 h) (n=4). BMDMs were either stimulated in suramin-containing media (no wash), washed 3x before stimulation (washout), or washed 3x with re-addition of each respective drug (washout + drug). **(F)** LDH and **(H)** IL-1β release in the supernatant from LPS-primed (1 μg mL^-1^, 4 h) human monocyte-derived macrophages (hMDMs) pre-treated with suramin (10 μM), 4-sulfonic calix[8]arene (10 μM), MCC950 (10 μM) or vehicle control (DMSO) before transfection with poly dA:dT (1 μg mL^-1^, 18 h) (n=4). **(H-I)** Electrostatic potential maps for docked suramin in AIM2 HIN domain (PDB:3RN5) using Autodock Vina. **(H)** Suramin conformer (conf1) identified from the pharmacophore model (docking of the 4-sulfonic calixarenes with AIM2) in the AIM2 HIN domain (monomer), **(I)** a possible alternative binding mode of co-crystal structure of suramin (grey) (from PDB:2NYR), (conf2, extended form), when docked between the 2 HIN domains of AIM2 (PDB:3RN5) (dimer). For electrostatic potential maps: blue: positive region, red: negative region, white: neutral region, red spheres: negatively charged atoms for the sulfonates of suramin. Images created using Pymol. Dose-response curves were fitted using a four parameter logistical (4PL) model. *p<0.05, **p<0.01, ****p>0.0001 determined by a two-way ANOVA with Sidak’s (E) or Dunnett’s (F,G) post hoc analysis. Values shown are mean ± the SEM.

Molecular modelling was used to assess if suramin could also interact with the dsDNA binding site of the HIN domain of AIM2. Suramin is a large polyanionic symmetrical urea molecule which is very flexible and therefore can adopt thousands of conformations (Fig 6A). The eleven conformers of suramin found in protein crystal structures from the Protein Data Bank (Supplementary Fig 7, Supplementary Table 4, (*41*–*49*)) were rigidly docked in the human AIM2 HIN domain (monomer) (PDB:3RN5)(*24*) using Autodock Vina(*26*). There were no favourable docking poses observed (Supplementary Table 5), therefore attention turned to finding alternative conformations of suramin. A conformational search for the ionised form of suramin at pH 7.4 was carried out in MOE (MOE 2015.08), with the 28 lowest energy conformations reported in Supplementary Table 6, however again none of these bound effectively to the AIM2 HIN domain (monomer) (Supplementary Table 7). Alternative conformations of suramin were then sought utilising the trigonal planar shaped pharmacophore model from the docking of the 4-sulfonic calixarenes with AIM2 (Supplementary Fig 8). Only one conformer of suramin (Fig 6H, trans cis, conf1) obtained from the pharmacophore-model showed good docking to the AIM2 HIN monomer, with 5 ionic interactions between the sulfonate groups and the HIN monomer domain residues (Fig 6I). The docked conformation also has two intramolecular hydrogen bonds forming 2 pseudo rings, 2 H-π interactions between one phenyl and the naphthalene to amino acid residues in the pocket and intramolecular π-π stacking interactions (Fig 6H, Supplementary Table 8). The largest distance measured between the sulfonates in the docked pose of suramin in 1 HIN domain was 17.6 Å close to that between the phosphates of dsDNA in the co-crystallized structure with a distance of 20.4 Å in the minor groove (Supplementary Fig 9A-B). The eleven suramin conformers from the crystal structures (*41*–*49*) were also docked across the two HIN domains (dimer) of AIM2 (PDB:3RN5)(*24*) (Supplementary Table 5), with potential binding spanning the major and minor grooves (Fig 6I). Only 1 conformer of suramin (co-crystal structure PDB:2NYR)(*41*) spanned the dimer (Fig 6I), forming interactions with both HIN domains with an O-O distance of 34.6 Å, similar to that observed for dsDNA with a distance of 34.7 Å (Supplementary Fig 9C). A good docking score of −22.0 Kcal/mol supported a strong binding interaction with suramin spanning across 2 HIN domains.

Molecular modelling was also used to assess whether 4-sulfonic calixarenes and suramin could also interact with cGAS, using the crystal structure of murine cGAS (PDB: 4LEZ, (*25*), resolution 2.30 Å). cGAS has a similar dsDNA interface to AIM2, made up of positively charged lysine and arginine residues, in addition to asparagine and histidine residues (Supplementary Fig 10A)(*24*). The molecular docking was validated by redocking the dsDNA ribose and phosphate backbone rigidly in the minor groove of cGAS using Autodock Vina(*26*), showing a good overlay between redocked dsDNA and co-crystallized dsDNA, with similar ionic interactions seen between the phosphate anions and the amino acid side chains (within 4.0Å) (Supplementary Fig 10A,B). Docking of 4-sulfonic calix[8]arene and suramin in cGAS in the dsDNA interface was carried out using MOE (MOE 2015.08) (flexible docking) and Autodock Vina (rigid docking). The preferred docked pose of 4-sulfonic calix[8]arene showed that the sulfonate groups bind in discrete regions mimicking dsDNA interactions with the positively charged amino acids (Supplementary Fig 10C). Similar distances were observed for the ionic interactions made by 4-sulfonic calix[8]arene, compared to the co-crystal structure with dsDNA (Supplementary Table 9). Docking of suramin in the minor groove region of cGAS agreed with the preliminary published study(*38*). The preferred docked conformers for suramin in cGAS form an increased number of ionic interactions between the sulfonate oxygen atoms and positively charged amino acid side chains, mimicking that of bound dsDNA. The binding modes of suramin in cGAS are similar to AIM2, with one U-shaped structure conformer (Conf3) (Supplementary Fig 10D) and one extended conformer (Conf4) (Supplementary Fig 10E). Both conformers bind to cGAS with good docking scores of ~−11 Kcal/mol (Supplementary Table 9), suggesting competitive binding with dsDNA.

## Discussion

dsDNA-driven AIM2 inflammasome responses are emerging as critical responses in the worsening of multiple diseases including atherosclerosis(*6*), cancer(*7*, *50*), ischemic stroke(*8*), and post-stroke immunosuppression(*10*). However, the tools to target and study AIM2 inflammasome responses are lacking, limited to knockdown/knockout of AIM2, non-selective suppressive oligonucleotides (such as A151(*51*)), or removal of extracellular dsDNA using DNases(*7*, *10*). Here we report the identification and characterisation of 4-sulfonic calixarenes as new and effective compounds at blocking dsDNA-driven inflammatory signalling, exhibiting high potency at the AIM2 receptor. We propose that 4-sulfonic calixarenes reversibly interact with the dsDNA binding site of AIM2 through exposed sulfonic acid groups, preventing binding of dsDNA and limiting AIM2 inflammasome formation.

The ability to limit AIM2-dependent inflammation could be beneficial in a range of disorders. Targeting IL-1β signalling reduces cardiovascular risk in atherosclerotic patients(*52*) and AIM2 has recently been identified as an essential mediator of atherosclerosis driven by clonal haematopoiesis induced by the common *Jak2^V617F^* mutation(*6*, *53*). AIM2 impedes anti-tumour responses in melanoma by limiting immunogenic type I IFN responses(*7*). AIM2 drives inflammation to worsen ischemic brain injury(*8*, *9*, *54*) and mediates post-stroke immunosuppression by induction of T cell death(*10*). AIM2 inflammasomes are also activated during radiotherapy, exacerbating radiation-induced tissue damage, and contributing to post-radiation side effects including immunosuppression, multi-organ dysfunction, and haemorrhage(*55*–*57*), although recent studies suggest AIM2 also contributes to the anti-tumour effects of radiotherapy(*58*). The discovery of suramin, a pre-existing clinically used drug, as an AIM2 and cGAS inhibitor will provide new opportunities to target diseases driven by dsDNA. Interestingly, suramin reduces brain injury following ischemic stroke(*59*), melanoma growth(*60*, *61*), and atherosclerosis(*62*), in animal models, but since these studies pre-date the identification of suramin as an inhibitor of dsDNA inflammatory signalling, it is unknown if suramin is exerting protection from these diseases via AIM2 inhibition. One consideration of using 4-sulfonic calixarenes or suramin as molecules to inhibit AIM2 *in vivo* is that these compounds are unlikely to pass the intact blood brain barrier due to the negative charge of the multiple sulfonate groups. Pharmacokinetic studies have shown that suramin is largely excluded from the brain(*63*). Further, following intravenous administration of a single dose of 4-sulfonic calix[4]arene, it is distributed widely in most organs, except the brain, is not metabolised, and is quickly cleared via the urine(*64*). However, as an approach to target dsDNA-induced inflammation in the brain post stroke, these molecules could still be effective in brain tissue due to the breakdown of the blood brain barrier following brain injury(*65*). Importantly, suramin and 4-sulfonic calixarenes also do not need to enter the brain to limit DNA-induced inflammasome activation systemically, such as immunosuppression post-stroke(*10*).

The inhibitory actions of 4-sulfonic calix[6]arene, 4-sulfonic calix[8]arene, and suramin on dsDNA-driven inflammation are not limited to AIM2 and these molecules are also effective at blocking cGAS- and TLR9-dependent inflammation. However, 4-sulfonic calix[6]arene and 4-sulfonic calix[8]arene exhibit enhanced potency for AIM2 compared to cGAS and TLR9 (IC_50_ values; 4-sulfonic calix[6]arene: AIM2 1.82 μM, cGAS 6.04 μM, TLR9 >100 μM; 4-sulfonic calix[8]arene: AIM2 0.21 μM, cGAS: 0.90 μM, TLR9: 14.26 μM) suggesting an increased preference for targeting AIM2 receptors. This increased preference for AIM2 was absent for suramin (IC_50_ values, AIM2: 1.48 μM, cGAS: 1.32 μM, TLR9: 31.2 μM), despite a similar proposed mechanism of inhibition. This suggests that the structure and arrangement of the sulfonic acid groups could be refined in future to produce molecules with enhanced specificity and potency for particular dsDNA sensors. Further, the dsDNA binding pockets of dsDNA binding proteins, such as AIM2 (PDB:3RN5)(*24*) and cGAS (PDB: 4LEZ)(*25*), has been resolved using cryo-EM to a high resolution, assisting with the generation of molecules highly specific to the dsDNA binding site of AIM2.

On the other hand, the ability of 4-sulfonic calixarenes and suramin to target multiple dsDNA PRRs may be advantageous in treating some diseases driven by dsDNA. The dsDNA inflammasome in humans is dependent on, at least in part, a cGAS-STING-NLRP3 pathway(*16*), suggesting that inhibition of both cGAS and AIM2 is required to fully prevent dsDNA-induced inflammasome responses in human. Several conditions have been reported to be worsened by both AIM2 inflammasomes and type I IFN signalling, such as ischaemic stroke(*66*) and atherosclerosis(*6*, *67*). Furthermore, genetic deletion of AIM2 leads to enhanced activation of the cGAS-STING pathway leading to increased type I IFN signalling(*68*–*70*). AIM2 depletion worsens outcomes in murine models of experimental autoimmune encephalomyelitis driven by enhanced type I IFN signalling(*71*, *72*). Interestingly, increased cGAS-STING signalling is beneficial in certain pathologies, such as cancer, where AIM2 depletion promotes anti-tumour responses by enhancing STING-dependent type I IFN responses(*7*), suggesting specific AIM2 inhibitors are useful but require careful consideration.

In summary, we have identified 4-sulfonic calix[6]arene, 4-sulfonic calix[8]arene, and suramin as inhibitors of dsDNA-driven inflammatory responses, providing new tools for the understanding of AIM2 and dsDNA responses in the development of disease. We have identified the mechanism of inhibition of these compounds, mediated by sulfonic acid groups that prevent dsDNA binding, assisting with the future drug development for more potent and specific inhibitors of AIM2, cGAS and TLR9.

## Acknowledgements

This study was supported by the Medical Research Council (MRC, UK) grant MR/T016515/1 (DB), the University of Manchester (Presidential Fellowship to JG), Yasser El-Sharkawy and a University of Manchester Presidential Scholars Award (LS), the European Research Council (ERC-StGs 802305 to AL), the German Research Foundation (DFG) under Germany’s Excellence Strategy (EXC 2145 SyNergy – ID 390857198) and by the DFG FOR 2879.

## Methods

### Reagents and Tools Table

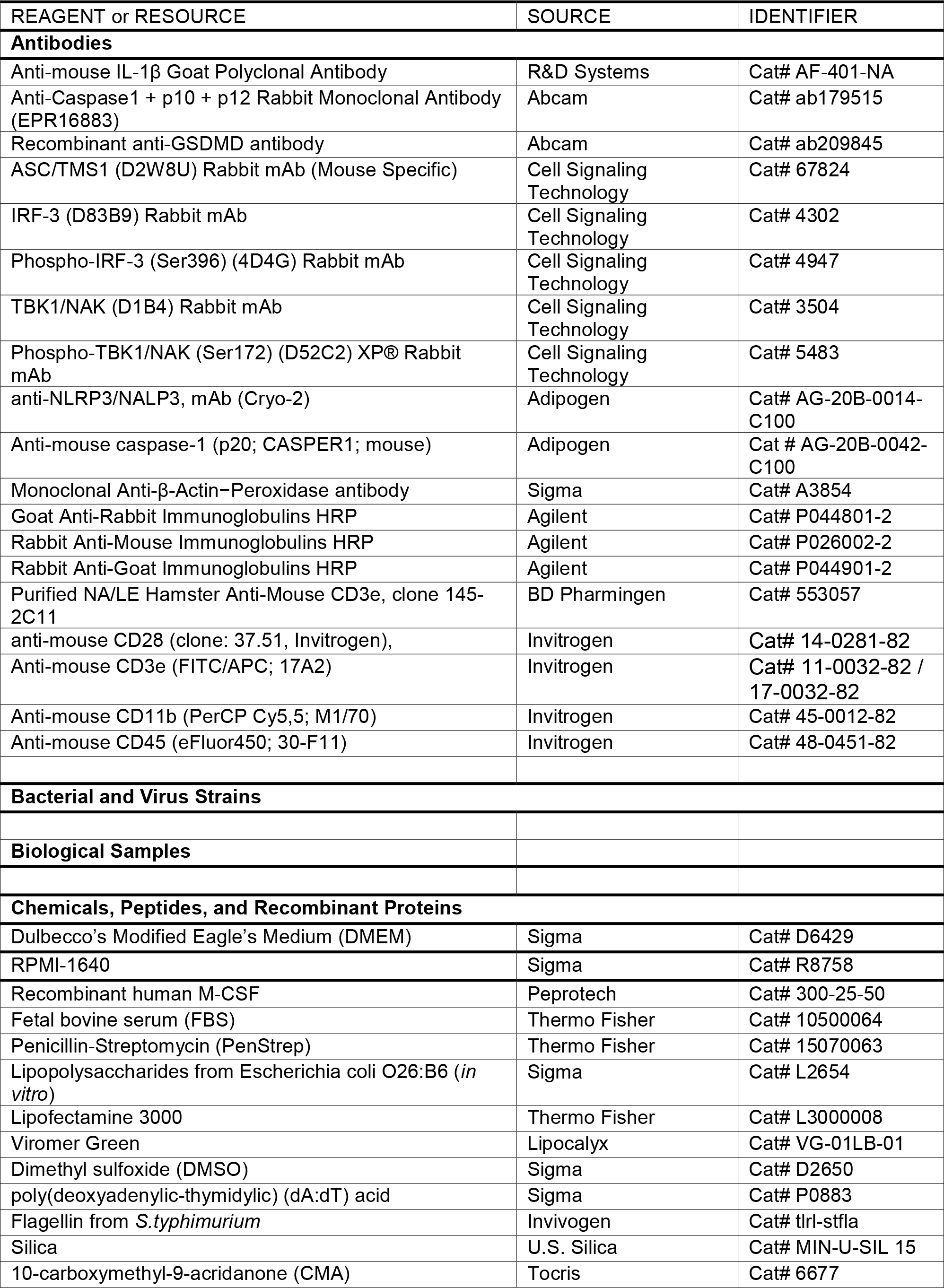
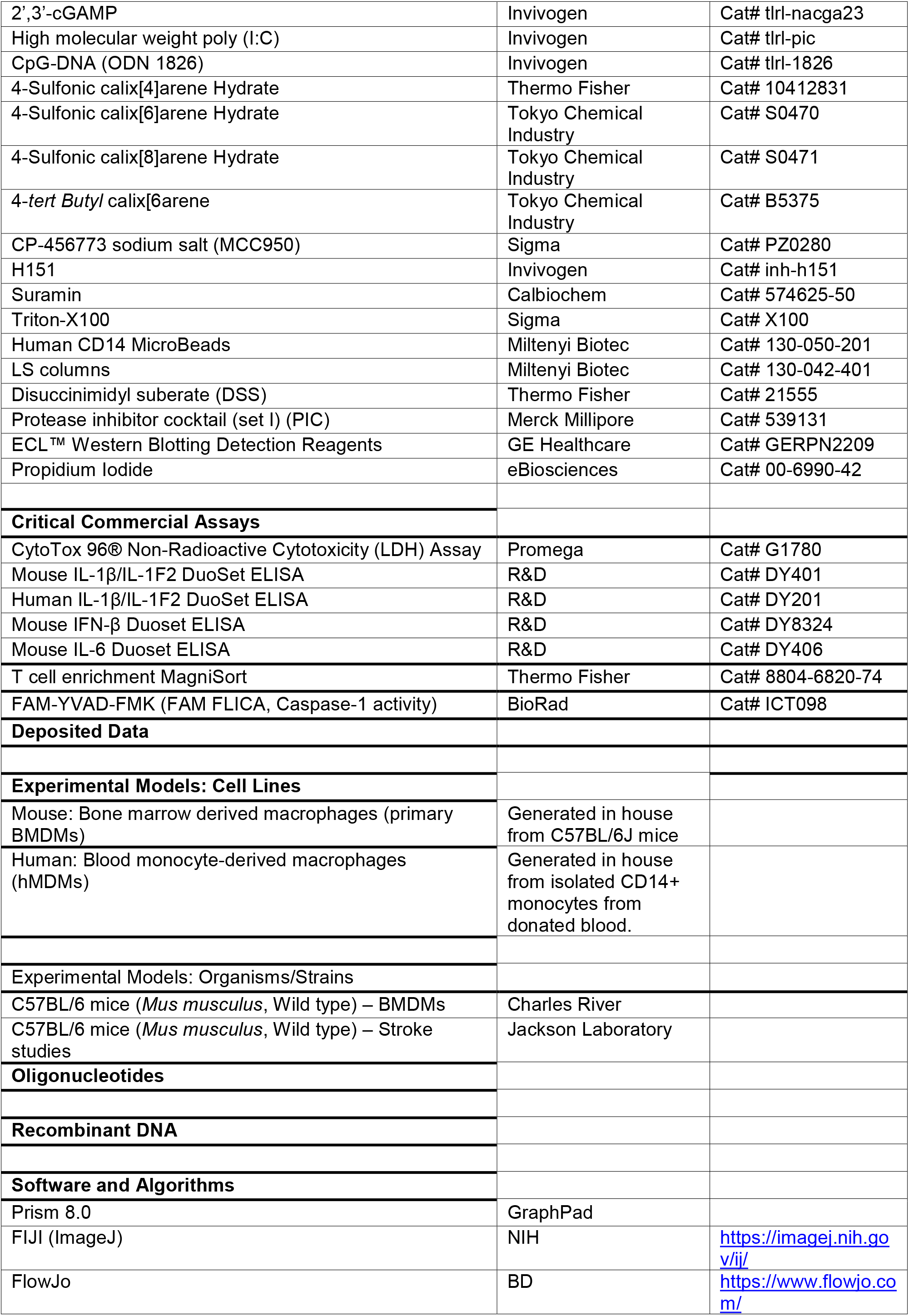
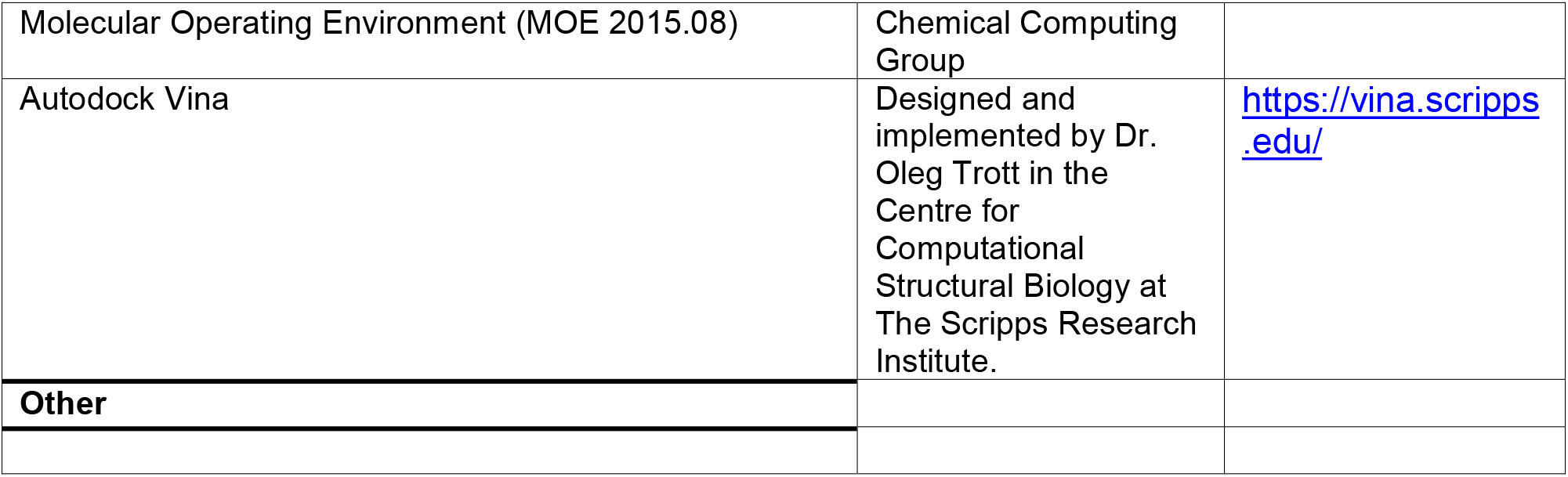

#### Cell culture

Primary bone marrow-derived macrophages (BMDMs) were isolated from male and female wild type C57BL/6J mice. All procedures were performed with appropriate personal and project licenses in place, in accordance with the Home Office (Animals) Scientific Procedures Act (1986) and approved by the Home Office and the local Animal Ethical Review Group, University of Manchester. Bone marrow from the femurs and tibias was collected, red blood cells were lysed by incubation in ACK lysis buffer, and the resulting marrow was cultured in 70% DMEM (10% v/v Foetal bovine serum (FBS), 100 U mL^-1^ penicillin, 100 μg mL^-1^ streptomycin) supplemented with 30% L929 conditioned media for 6-7 days. BMDM cultures were fed with media (containing 30% v/v L929 conditioned media) on day 3. Before experiments, BMDMs were scraped and seeded out at a density of 1 x 10^6^ mL^-1^ in DMEM (10% v/v FBS, 100 U mL^-1^ penicillin, 100 μg mL^-1^ streptomycin) overnight.

Human monocyte-derived macrophages (hMDMs) were differentiated from isolated CD14^+^ monocytes. Human peripheral blood mononuclear cells (PBMCs) were isolated from leucocyte cones obtained from the National Blood Transfusion Service (Manchester, UK) with full ethical approval from the research governance, ethics, and integrity committee at the University of Manchester (ref. 2018-2696-5711). PBMCs were collected by density centrifugation using a 30% Ficoll gradient and monocytes were positively selected using magnetic CD14^+^ microbeads and LS columns (Miltenyi). hMDMs were generated by culturing CD14^+^ monocytes in RPMI-1640 (10% v/v FBS, 100 U mL^-1^ penicillin, 100 μg mL^-1^ streptomycin, 2 mM L-glutamine) supplemented with 50 ng mL^-1^ recombinant human M-CSF for 7 days. hMDMs cultures were fed with media (containing 50 ng mL^-1^ M-CSF) on day 3. Before experiments hMDMs were scraped and seeded at a density of 1 x 10^6^ mL^-1^ in RPMI-1640 (10% v/v FBS, 100 U mL^-1^ penicillin, 100 μg mL^-1^ streptomycin, 2 mM L-glutamine) overnight.

#### Inflammasome activation assays

Murine BMDMs were LPS-primed (1 μg mL^-1^, 4 hours) in DMEM (10% v/v FBS, 100 U mL^-1^ penicillin, 100 μg mL^-1^ streptomycin). After LPS-priming, BMDMs were incubated in serum free DMEM and the inflammasome was activated by lipofectamine 3000-mediated transfection of poly dA:dT (1 μg mL^-1^, 4 hours) for AIM2, or bacterial flagellin (1 μg mL^-1^, 4 hours) for NLRC4, or by addition of silica crystals (300 μg mL^-1^) to activate NLRP3. hMDMs were LPS-primed (1 μg mL^-1^, 4 hours) in RPMI-1640 (10% v/v FBS, 100 U mL^-1^ penicillin, 100 μg mL^-1^ streptomycin, 2 mM L-glutamine). After LPS-priming, hMDMs were incubated in RPMI 1640 (1% v/v FBS) and the inflammasome was activated by Viromer Green-mediated transfection of poly dA:dT (1 μg mL^-1^, 18 hours). Control experiments were performed using transfection reagent alone in all experiments (mock). When used 4-sulfonic calixarenes, MCC950, H151 or suramin were incubated with the BMDMs at the indicated concentrations in serum free DMEM for 15 minutes before stimulation with inflammasome activators.

Cell supernatants were collected for analysis of IL-1β release and pyroptosis. IL-1β release was determined by ELISA (R&D systems) and pyroptosis was determined by lactate dehydrogenase (LDH) release (Promega) according to the manufacturer’s instructions.

#### cGAS-STING activation assays

For the detection of interferon (IFN)-β release in the supernatant, murine BMDMs were seeded into 96 well plates, incubated in serum free DMEM and transfected using lipofectamine 3000 with poly dA:dT (1 μg mL^-1^), 2’,3’-cGAMP (1.5 μg mL^-1^) or treated with CMA (250 μg mL^-1^) for 6 hours. The supernatant was collected and analysed for IFN-β release by ELISA (R&D systems). For the detection of phosphorylated IRF-3 (S396) and TBK-1 (S172), BMDMs were seeded into 24 well plates, incubated in serum free DMEM and transfected using lipofectamine 3000 with poly dA:dT (1μg mL^-1^) for 3 hours. Following stimulation, the supernatant was discarded and cells were lysed with Triton lysis buffer (50 mM Tris-HCl, 150 mM NaCl, 1% (v/v) Triton x-100, protease and phosphatase inhibitor cocktail) and analysed by western blot. Control experiments were performed using transfection reagent alone in all experiments (mock). When used 4-sulfonic calixarenes, H151 or suramin were incubated with the BMDMs at the indicated concentrations in serum free DMEM for 15 minutes before stimulation with inflammasome activators.

#### TLR activation assays

Murine BMDMs in 96 well plates were incubated in serum DMEM with the indicated concentration of 4-sulfonic calixarene for 15 minutes, before stimulation with either the TLR9 agonist CpG-DNA (1 μM), the TLR4 agonist LPS (1 μg mL^-1^) or the TLR3 agonist Poly I:C (5 μg mL^-1^) for 6 hours. Supernatants were analysed for cytokine release using IL-6 ELISA (R&D systems) for CpG-DNA and LPS or IFN-β ELISA (R&D systems) for Poly I:C, performed according to the manufacturer’s instructions. Additionally, cells were lysed with Triton lysis buffer (50 mM Tris-HCl, 150 mM NaCl, 1% (v/v) Triton x-100, protease inhibitor cocktail) and analysed by western blot.

#### Washout experiments

Murine BMDMs were LPS-primed (1 μg mL^-1^, 4 hours) in DMEM (10% v/v FBS, 100 U mL^-1^ penicillin, 100 μg mL^-1^ streptomycin). After LPS-priming, BMDMs were incubated in serum-free DMEM with DMSO (0.5% v/v), 30 μM of 4-sulfonic calix[4]arene, 4-sulfonic calix[6]arene, 4-sulfonic calix[8]arene or 10 μM of suramin for 1 hour. BMDMs were washed 3x with warm SF DMEM (37 °C) and incubated at 37 °C for 5 minutes between washes. Following washes, the AIM2 inflammasome was activated as described above. In each experiment, controls were performed in parallel including BMDMs that were not washed, and BMDMs that had fresh drug added after the washes before stimulation. Supernatants were collected and analysed for inflammasome activation by IL-1β release (determined by IL-1β ELISA) and pyroptosis (determined by LDH release).

#### Western blot

Cell lysates were diluted with 5X Laemmli buffer and boiled (95 °C, 5 minutes) before being resolved by tris-glycine SDS-PAGE and transferred onto nitrocellulose or PVDF membranes at 25V using a semidry Trans-Blot Turbo (BioRad). Membranes were blocked for 1 hour with milk (5% w/v) in PBS-Tween (PBS-T, 0.1% v/v Tween 20) and overnight incubation with the indicated primary antibodies in BSA (5% w/v) PBS-T. Membranes were then incubated with the corresponding HRP-tagged secondary antibodies in BSA (5% w/v) PBS-T for 1 hour before visualisation using Amersham ECL prime detection reagent (GE healthcare) and a G:Box Chemi XX6 system (Syngene). Western blots for phosphorylated proteins used Tris-buffered saline (TBS) instead of PBS. Densitometry was performed using Fiji (ImageJ).

#### ASC oligomerisation

For western blotting of insoluble ASC oligomers and markers of inflammasome activation, 1 x 10^6^ BMDMs were seeded into 12 well plates and stimulated as described above to activate the inflammasomes. Total cell lysates (combined supernatant and cell lysate) were made by directly adding protease inhibitor cocktail followed by Triton x-100 (Tx-100, 1% v/v) to the supernatant of each well. BMDM lysates were separated into Tx-100 soluble and insoluble fractions by centrifugation at 6800*xg* for 20 minutes at 4°C. The resulting pellet (insoluble fraction) was resuspended in PBS with disuccinimidyl suberate (DSS, 2 mM, 30 min, RT) to chemically crosslink ASC oligomers. Following crosslinking, the Tx-100 insoluble fraction was spun at 6800*xg* for 20 minutes at 4°C and the resulting pellet was resuspended in 1X Laemmli buffer and boiled (95°C, 5 minutes). The Tx-100 soluble fraction was concentrated by trichloroacetic acid (TCA) precipitation. Proteins were precipitated by mixing TCA (20% w/v in ddH2O) at a 1:1 ratio with the Tx-100 soluble fraction and spun at 14000*xg* for 10 minutes. The precipitated protein pellet was washed in acetone, before an additional spin at 14000*xg* and resuspension and boiling (95°C, 5 minutes) in 2X Laemmli buffer.

##### In vitro T-cell death assays

###### Whole splenocyte culture

Whole splenocyte culture was performed as previously described (*10*). Spleens from naïve wild-type mice (C57BL/6J) were dissected and single splenocyte suspension prepared by mincing and using a 40 μm cell strainer. Cells were washed with PBS, cell numbers and viability was assessed using an automated cell counter (Biorad, Germany). The required viability threshold was ≥ 80 %. Cells were then cultured (RPMI 1640, 10% (v/v) heat-inactivated FBS, 1 % (v/v) penicillin/streptomycin, 10 μM β-mercaptoethanol) on a 96 well flat bottom plate at a density of 100,000 cells per well in a final volume of 200 μL. Prior to the stimulation, cells were stimulated with serum from either stroke or sham operated mice at a concentration of 25 % (v/v) total well volume for 16 h. After stimulation, cell numbers were analysed by flow cytometry using antibodies against CD45, CD3 and CD11b.

###### T cell isolation and culture

Round-bottom 96-well plates were coated with 100 μL of PBS containing a mixture of 0.5 mg mL^-1^ purified NA/LE hamster anti-mouse CD3e (clone: 145-2C11, BD Pharmingen) and 0.5 mg mL^-1^ anti-mouse CD28 (clone: 37.51, Invitrogen), and then incubated overnight. Spleens, isolated from mice (C57BL/6J), were homogenized into single splenocyte suspensions by using a 40 μm cell strainer, erythrocytes were lysed using isotonic ammonium chloride buffer. T cells were purified from splenocytes using a negative selection kit (MagniSort T cell enrichment, Thermo Fisher) according to the manufacturer’s instructions. Purity was reliably ≥ 90 % as assessed by flow cytometry. Cells were resuspended in complete RPMI 1640 (GIBCO) and supplemented with 10 % (v/v) FBS, 1 % (v/v) penicillin/streptomycin and 10 μM β-mercaptoethanol. T cells were seeded into the anti-CD3/CD28 coated plates at a density of 300,000 cells per well in a total volume of 200 μL.

###### BMDM T cell co-culture

BMDM T cell co-culture was performed as previously described (*73*). BMDMs were cultured for 7 days, washed, harvested, counted, and seeded in flat bottom tissue culture-treated 96 well plate at a density of 100,000 cells per well in a final volume of 200 μL. Cells were cultured overnight. BMDMs were then primed for 4 h with LPS (100 ng mL^-1^). For the last hour of LPS priming 4-sulfonic calix[6]arene was added in a final concentration of 10 μM to the BMDMs. Then BMDMs were carefully washed and afterwards stimulated for 10 minutes with serum from either stroke or sham-operated C57BL6/J mice at a concentration of 25% (v/v) total volume. Control-treated BMDMs received only FBS-containing culture media. After stimulation, the culture medium was removed, and the cells were washed with sterile PBS to ensure no leftover serum in the medium. BMDM-T cell interaction via cell-cell contact was then assessed. T cells were added to the serum-stimulated BMDMs at a density of 200,000 cells per well in a total volume of 200 μL complete RPMI medium (10 % (v/v) FBS, 1 % (v/v) penicillin/streptomycin and 10 μM β-mercaptoethanol), and then incubated for 4h at 37 °C with 5 % CO2. T cell counts and survival rate were assessed by flow cytometry. 240 minutes after addition of T cells to the serum-stimulated BMDMs, cells were harvested and analyzed. For the analysis of T cell death, we analyzed Propidium Iodide (PI) uptake by T cells after addition of PI in a final concentration of 1 mg mL^-1^ to the co-culture medium via flow cytometry. The number of PI^+^ T cells was quantified and normalized to the corresponding control group (sham serum treated).

###### Animal experiments

All animal experiments were performed in accordance with the guidelines for the use of experimental animals and were approved by the governmental committees (Regierungspräsidum Oberbayern). Wildtype C57BL/6J mice were obtained from Jackson Laboratory (Bar Harbor, USA). All mice were housed with free access to food and water at a 12 h dark-light cycle. For this exploratory study, animal numbers were estimated based on previous results from transient ischemia-reperfusion stroke model on extent and variability of T cell death after stroke. Data were excluded from all mice that died during surgery. Detailed exclusion criteria are described below. Animals were randomly assigned to treatment groups and all analyses were performed by investigators blinded to group allocation. All animal experiments were performed and reported in the accordance with the ARRIVE guidelines.

###### Induction of Ischaemic stroke and T-cell death in vivo

Transient ischemia-reperfusion stroke model. Mice were anaesthetized with isoflurane delivered in a mixture of 30 % O2 and 70 % N2O. An incision was made between the ear and the eye in order to expose the temporal bone. Mice were placed in supine position, and a laser Doppler probe was affixed to the skull above the middle cerebral artery (MCA) territory. The common carotid artery and left external carotid artery were exposed via midline incision and further isolated and ligated. A 2-mm silicon-coated filament (Doccol) was inserted into the internal carotid artery, advanced gently to the MCA until resistance was felt, and occlusion was confirmed by a corresponding decrease in blood flow (i.e., a decrease in the laser Doppler flow signal by ≥ 80 %. After 60 minutes of occlusion, the animals were re-anesthetized, and the filament was removed. After recovery, the mice were kept in their home cage with ad libitum access to water and food. Sham-operated mice received the same surgical procedure, but the filament was removed in lieu of being advanced to the MCA. Body temperature was maintained at 37°C throughout surgery in all mice via feedback-controlled heating pad. The overall mortality rate of animals subjected to MCA occlusion was approximately 20%. All animals in the sham group survived the procedure. Exclusion criteria: 1. Insufficient MCA occlusion (a reduction in blood flow to > 20 % of the baseline value). 2. Death during the surgery. 3. Lack of brain ischemia as quantified post-mortem by histological analysis.

###### Organ and tissue processing

Mice were deeply anaesthetized with ketamine (120 mg kg^-1^) and xylazine (16 mg kg^-1^) and blood was drawn intracardially into low-bind collection tubes. Blood was incubated for 10 minutes at room temperature, centrifuged for 10 minutes at 3,000x*g*, serum collected and stored at 80°C until further use.

Spleen was transferred to tubes containing Hank’s balanced salt solution (HBSS), homogenized and filtered through 40 μm cell strainers to obtain single cell suspensions. Homogenized spleens were subjected to erythrolysis using isotonic ammonium chloride buffer.

###### Splenic T cell analysis via fluorescence-activated Cell Sorting (FACS)

The anti-mouse antibodies listed below were used for surface marker staining of CD45^+^ leukocytes, CD45^+^CD11b^+^ monocytes and CD3^+^ T cells. Fc blocking (Anti CD16/CD32, Invitrogen, US) was performed on all samples prior to extracellular antibody staining. All stains were performed according to the manufacturer’s protocols. Flow cytometric data were acquired using a Cytek Northern Light flow cytometer (Cytek, US) and analyzed using FlowJo software (Treestar, US).

###### FAM660 caspase-1 staining for flow cytometry

To detect the active forms of caspase-1 in spleen, cell suspensions were stained with the fluorescent inhibitor probe FAM-YVAD-FMK (FAM 660, BioRad, Germany) for 30 minutes at 37°C according to the manufacturer’s instructions. After washing, the cells were stained for CD45^+^CD3^+^ T cells and CD45^+^CD11b^+^ monocytes. The flow cytometry data were acquired on a Cytek Northern Light flow cytometer (Cytek, US).

###### Conformational search

Conformational searches were carried out in Molecular Operating Environment (MOE 2015.08, Chemical Computing Group, Canada) for the ionized forms of 4-sulfonic calix(X)arenes and suramin at physiological pH (7.4) using the stochastic search method. A rejection limit of 100 was chosen and iteration limit of 10,000. RMS gradient was set to 0.01 kcal mol^-1^ with a MM iteration limit of 500. The RMSD limit was set to 0.15 Å with a defined energy window of 7 kcal mol^-1^ and conformation limit of 10,000 conformers. Energy values in Kcal mol^-1^ were ranked based on ascending order and the top 50, 75 and 100 were chosen for the 4, 6 and 8 membered calixarenes, respectively, with the top 49 conformations reported in Supplementary Table 1. A stochastic search was carried out for low energy conformers of the *trans cis, trans trans* and *cis cis* urea conformations of suramin. The lowest energy conformers from each were chosen to undergo rigid docking using Autodock Vina (*26*).

###### Docking validation

The dsDNA was removed from the protein prior to docking. Polar hydrogens were added as well as Gasteiger charges. dsDNA ribose and phosphate backbone were redocked rigidly in Autodock Vina (*26*) between 2 adjacent HIN domains. AIM2 HIN domain (PDB:3RN5) (*24*). A grid box was created with x, y, z dimensions of 48 Å, 52 Å and 36 Å, and centered with x, y, z coordinates of 23.636 Å, 18.027 Å and 22.444 Å. cGAS monomer (PDB:4LEZ) (*25*). A grid box was created with x, y, z dimensions of 30 Å, 58 Å and 56 Å, and centered with x, y, z coordinates of −27.364 Å, −30.065 Å and 12.776 Å. Two software packages were used for molecular validation of the docked inhibitors which were MOE and Autodock Vina for rigid docking.

###### Docking of 4-sulfocalixarenes and suramin with AIM2 and cGAS

The DNA was removed from the protein before docking the lowest energy conformer for each of the different conformer types of 4-sulfonic calix[4]arene, 4-sulfonic calix[6]arene and 4-sulfonic calix[8]arene, and the 28 generated conformers of suramin obtained from MOE. The AIM2 protein (PDB:3RN5) (*24*) was already solvated and the bound DNA was removed prior to docking. Protonation and protein structural preparation, including tautomerization states of histidine residues, was carried out. 10-20 docking poses were set for each using triangle matcher as the placement method (for 4-sulfonic calixarenes) and pharmacophore placement (for suramin docking in the HIN domain monomer) using London dG as a scoring function and rescoring was carried out. Docking was carried out between two HIN domains for suramin using the same set of parameters as that used for the 4-sulfocalixarene docking, with amino acid residues chosen spanning the interface of the major groove region of the dsDNA. Molecular docking of the 4-sulfocalixarenes and suramin conformers in AIM2 using AutoDock Vina (PDB:3RN5) (*24*) was carried out in the monomer. A grid box was created for docking in the monomeric HIN domain with x, y, z dimensions of 52 Å, 46 Å and 30 Å, and centered with x, y, z coordinates of 17.893 Å, 6.281 Å and 9.642 Å. Rigid docking for 20 GA runs was carried out for 4-sulfocalixarenes and suramin. Suramin was redocked also using the grid box created for redocking of dsDNA between 2 HIN domains to provide a potential alternative binding mode. 4-Sulfocalix[8]arene and suramin were docked using MOE. Molecular validation was carried out by redocking in Autodock Vina using the same grid box created for redocking dsDNA (ribose and phosphate backbone) in cGAS (PDB:4LEZ) (*25*).

###### Quantification and statistical analysis

Data are presented as mean values plus the SEM. Accepted levels of significance were *p<0.05, **p<0.01, ***p<0.001, ****p<0.0001. Statistical analyses were carried out using GraphPad Prism (version 8). Equal variance and normality were assessed with Levene’s test and the Shapiro–Wilk test, respectively, and appropriate transformations were applied or analysed using Mann-Whitney U test or Kruskal-Wallis test with Dunn’s post hoc analysis. Groups containing normally distributed data were analysed using a two-way ANOVA with Dunnett’s post hoc analysis. n represents experiments performed on individual animals or cells acquired from individual animals. Dose-response curves were fitted using a four parameter logistical (4PL) model.

