## Supplementary figures for "Discovery of an AIM2 inflammasome inhibitor for the treatment of DNA-driven inflammatory disease"

Supplementary Figure 1

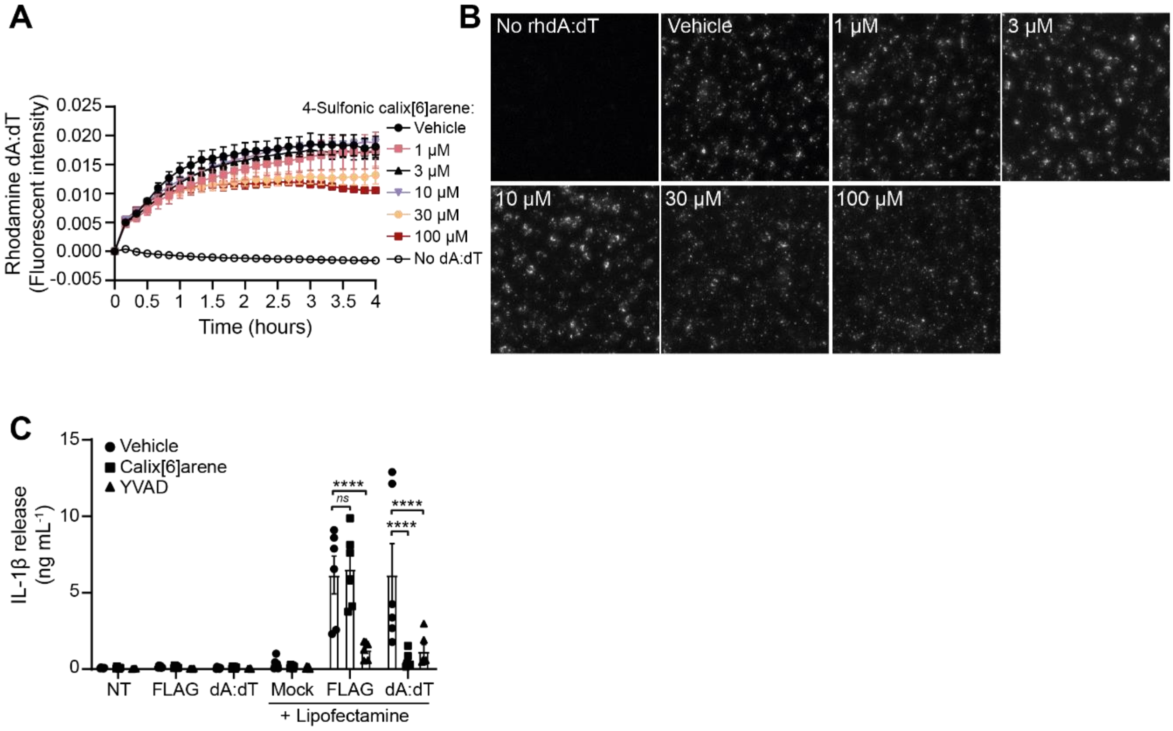


**Supplementary Figure 1 – 4-Sulfonic calix[6]arene does not impact transfection. (A)** Fluorescence intensity of rhodamine-tagged poly dA:dT (1 µg mL$^{-1}$) transfected into bone marrow-derived macrophages (BMDMs) in the presence of the indicated concentration of 4-sulfonic calix[6]arene (1-100 µM) (n=3). **(B)** Representative images of BMDMs from the experiment in (A). **(C)** IL-1β release in the supernatants of LPS-primed (1 µg mL$^{-1}$, 4 h) BMDMs. BMDMs were pre-treated with 4-sulfonic calix[6]arene (30 µM), the caspase-1 inhibitor Ac-YVAD-CMK (YVAD, 100 µM) or vehicle control (DMSO) before stimulation with poly dA:dT (1 µg mL$^{-1}$) or flagellin (1 µg mL$^{-1}$) in the presence or absence of lipofectamine 3000 for 4 h (n=6). **p<0.01, ****p>0.0001 determined by a two-way ANOVA with Dunnett's post hoc analysis vs vehicle control. Values shown are mean ± the SEM.

Supplementary Figure 2

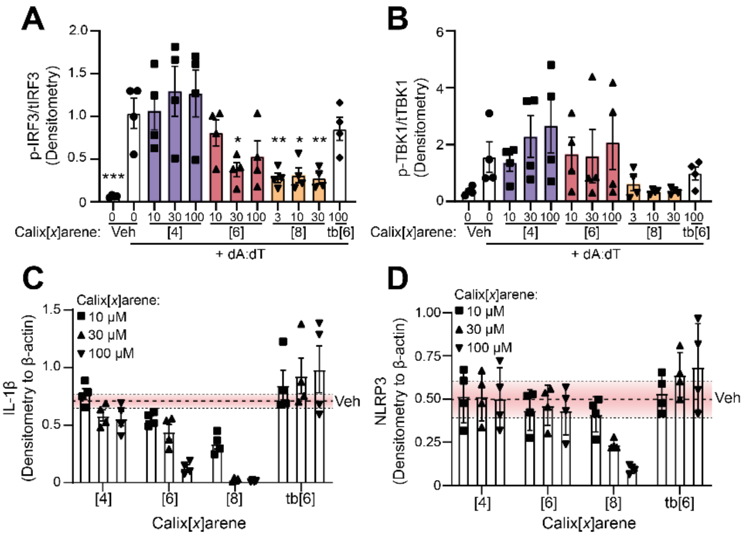

**Supplementary Figure 2 – Densitometry from Figure 3: Inhibition of dsDNA inflammatory signalling by 4-sulfonic calixarenes is readily reversible and is mediated by the exposed sulfonic acid groups. (A)** The ratio of p-IRF3 (Ser396) to total IRF3 and **(B)** the ratio of p-TBK1 to total TBK1 determined by densitometry of experiments shown in Fig 3G (n=4). **(C)** Densitometry of IL-1β and **(D)** NLRP3 from experiments shown in Fig 3I (n=4). *p<0.05, **p<0.01, ***p>0.001 determined by a one-way ANOVA with Dunnett's post hoc analysis vs vehicle control. Values shown are mean ± the SEM.

Supplementary Figure 3

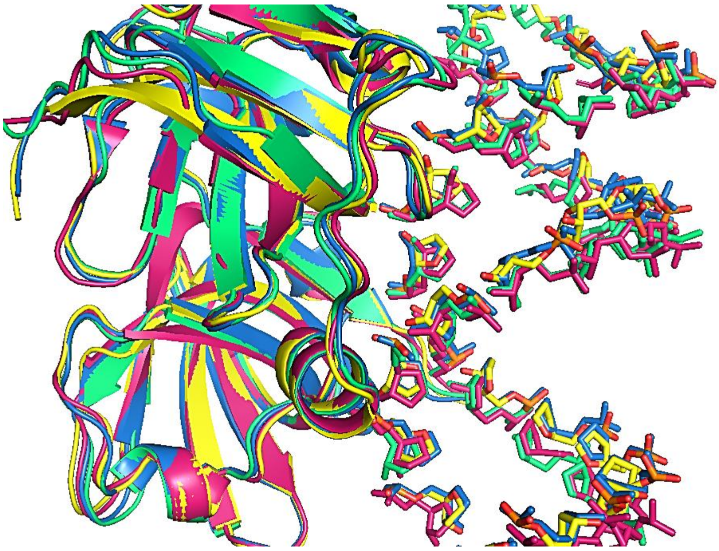

**Supplementary Figure 3 -** Superimposition of the 4 HIN domains of AIM2 with dsDNA phosphate and ribose backbone from the crystal structure bound to each showing binding of phosphates in discrete regions, blue: (HIN-a), green (HIN-b), yellow (HIN-c), red (HIN-d) (PDB: 3RN5). Image created using Pymol.

Supplementary figure 4

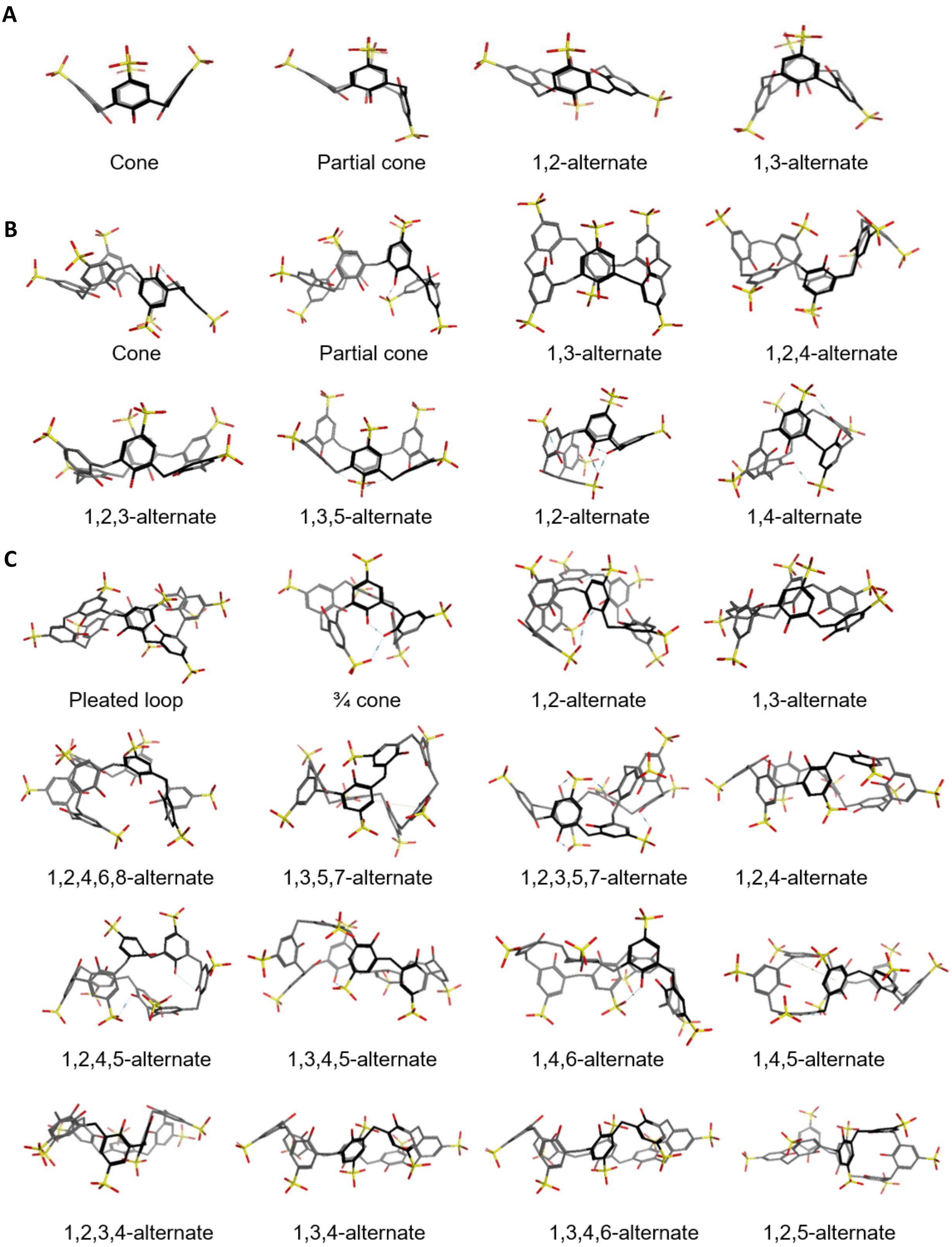


**Supplementary Figure 4 -** Stick diagrams for the different conformers of **(A)** 4-sulfonic calix[4]arene, **(B)** 4-sulfonic calix[6]arene and **(C)** 4-sulfonic calix[8]arene using MOE.

**Supplementary Table 1 -** Top ranked 4-sulfonic calixarene conformers and their energies obtained from a stochastic search in MOE.

| | 4-Sulfonic calix[4]arene | | | 4-Sulfonic calix[6]arene | | | 4-Sulfonic calix[8]arene | | |
|---|---|---|---|---|---|---|---|---|---|
| **#** | **Conformation** | **E (Kcal/mol)** | **ΔE (Kcal/mol)** | **Conformation** | **E (Kcal/mol)** | **ΔE (Kcal/mol)** | **Conformation** | **E (Kcal/mol)** | **ΔE (Kcal/mol)** |
| **1** | partial cone | 238.50 | 0.00 | 1,2,4-alternate | 335.28 | 0.00 | 1,2,4,6,8-alternate | 431.59 | 0.00 |
| **2** | cone | 239.62 | +1.12 | 1,3,5-alternate | 335.30 | +0.02 | 1,2,4,6,8-alternate | 433.82 | +2.23 |
| **3** | 1,3-alternate | 241.41 | +2.91 | 1,4-alternate | 335.33 | +0.05 | 1,2-alternate | 436.07 | +4.48 |
| **4** | 1,2-alternate | 241.97 | +3.47 | 1,3,5-alternate | 335.36 | +0.08 | 1,2,4,6,8- alternate | 437.01 | +5.42 |
| **5** | 1,2-alternate | 241.97 | +3.47 | 1,3,5-alternate | 335.39 | +0.11 | 1,2,4,6,8- alternate | 437.49 | +5.90 |
| **6** | 1.2-alternate | 241.98 | +3.48 | 1,3,5-alternate | 335.44 | +0.16 | ¾ cone | 437.58 | +5.99 |
| **7** | 1,2-alternate | 241.99 | +3.49 | 1,3,5-alternate | 338.73 | +3.45 | 1,3,5,7-alternate | 438.14 | +6.55 |
| **8** | 1,3-alternate | 242.28 | +3.78 | 1,3,5-alternate | 338.77 | +3.49 | ¾ cone | 438.17 | +6.58 |
| **9** | 1,3-alternate | 242.28 | +3.78 | 1,3,5-alternate | 338.86 | +3.58 | ¾ cone | 438.37 | +6.78 |
| **10** | 1,2-alternate | 242.32 | +3.82 | 1,2,4-alternate | 338.86 | +3.58 | 1,2,4,6,8- alternate | 438.66 | +7.07 |
| **11** | 1,3-alternate | 242.35 | +3.85 | 1,2,4-alternate | 338.87 | +3.59 | 1,3,5,7-alternate | 439.17 | +7.58 |
| **12** | 1,3-alternate | 242.36 | +3.86 | 1,2,4-alternate | 338.90 | +3.62 | 1,2,4,6,8- alternate | 440.26 | +8.67 |
| **13** | 1,3-alternate | 242.37 | +3.87 | 1,4-alternate | 338.92 | +3.64 | 1,3,5,7-alternate | 440.44 | +8.85 |
| **14** | 1,3-alternate | 242.73 | +4.23 | 1,3,5-alternate | 338.93 | +3.65 | 1,2,4,6,8- alternate | 440.77 | +9.18 |
| **15** | partial cone | 242.78 | +4.28 | 1,3,5-alternate | 338.94 | +3.66 | 1,2,4,6,8- alternate | 440.80 | +9.21 |
| **16** | partial cone | 242.78 | +4.28 | 1,3,5-alternate | 338.94 | +3.66 | 1,3,5,7-alternate | 440.81 | +9.22 |
| **17** | 1,3-alternate | 242.78 | +4.28 | 1,3,5-alternate | 339.00 | +3.72 | 1,3,5,7-alternate | 440.82 | +9.23 |
| **18** | 1,3-alternate | 242.80 | +4.3 | 1,3,5-alternate | 339.00 | +3.72 | 1,2,3,4-alternate | 440.88 | +9.29 |
| **19** | 1,3-alternate | 242.81 | +4.31 | 1,3,5-alternate | 339.00 | +3.72 | 1,2,4,6,8- alternate | 440.89 | +9.30 |
| **20** | 1,3-alternate | 242.86 | +4.36 | 1,3,5-alternate | 339.01 | +3.73 | 1,3,5,7-alternate | 440.89 | +9.30 |
| **21** | 1,2-alternate | 242.88 | +4.38 | 1,3,5-alternate | 339.06 | +3.78 | 1,3,5,7-alternate | 441.11 | +9.52 |
| **22** | 1,3-alternate | 242.88 | +4.38 | 1,3,5-alternate | 339.07 | +3.79 | 1,2,4,6,8- alternate | 441.41 | +9.82 |
| **23** | partial cone | 242.89 | +4.39 | 1,3,5-alternate | 339.09 | +3.81 | 1,3,5,7-alternate | 441.422 | +9.832 |
| **24** | partial cone | 242.90 | +4.40 | 1,3,5-alternate | 339.12 | +3.84 | 1,3,5,7-alternate | 442.09 | +10.50 |
| **25** | partial cone | 242.90 | +4.40 | 1,2,4-alternate | 339.15 | +3.87 | 1,2,4,5-alternate | 442.16 | +10.57 |
| **26** | partial cone | 242.90 | +4.40 | 1,2,4-alternate | 339.16 | +3.88 | 1,2,4,6,8- alternate | 442.50 | +10.91 |
| **27** | partial cone | 242.90 | +4.40 | 1,2,4-alternate | 339.16 | +3.88 | 1,2,4,6,8- alternate | 442.57 | +10.98 |
| **28** | 1,3-alternate | 242.91 | +4.41 | 1,2,4-alternate | 339.16 | +3.88 | 1,2,4,6,8- alternate | 442.58 | +10.99 |
| **29** | partial cone | 243.26 | +4.76 | 1,2,4-alternate | 339.18 | +3.90 | 1,2,4,6,8- alternate | 442.64 | +11.05 |
| **30** | 1,3-alternate | 243.33 | +4.83 | 1,2,4-alternate | 339.23 | +3.95 | 1,2,4,6,8- alternate | 442.92 | +11.33 |
| **31** | partial cone | 243.34 | +4.84 | 1,3,5-alternate | 339.33 | +4.05 | Pleated loop | 443.15 | +11.56 |
| **32** | 1,3-alternate | 243.77 | +5.27 | 1,3,5-alternate | 339.33 | +4.05 | 1,2,4,6,8- alternate | 443.17 | +11.58 |
| **33** | cone | 243.80 | +5.30 | 1,2,4-alternate | 339.35 | +4.07 | 1,2,4,6,8- alternate | 443.20 | +11.61 |
| **34** | 1,3-alternate | 243.80 | +5.30 | 1,3,5-alternate | 339.39 | +4.11 | 1,2,3,5,7- alternate | 443.21 | +11.62 |
| **35** | 1,3-alternate | 243.85 | +5.35 | 1,3,5-alternate | 339.41 | +4.13 | 1,2,3,5,7- alternate | 443.21 | +11.62 |
| **36** | 1,3-alternate | 243.96 | +5.46 | 1,3,5-alternate | 339.41 | +4.13 | 1,2,3,5,7- alternate | 443.26 | +11.67 |
| **37** | partial cone | 243.98 | +5.48 | 1,3,5-alternate | 339.45 | +4.17 | 1,2,4,6,8- alternate | 443.33 | +11.74 |
| **38** | 1,3-alternate | 243.98 | +5.48 | 1,3,5-alternate | 339.61 | +4.33 | Pleated loop | 443.41 | +11.82 |
| **39** | partial cone | 244.24 | +5.74 | 1,3,5-alternate | 339.61 | +4.33 | 1,3,5,7-alternate | 443.45 | +11.86 |
| **40** | 1,3-alternate | 244.26 | +5.76 | 1,3,5-alternate | 339.76 | +4.48 | 1,2,4,6,8- alternate | 433.58 | +1.99 |
| **41** | partial cone | 244.30 | +5.80 | 1,2,4-alternate | 339.96 | +4.68 | 1,2,4,6,8- alternate | 443.61 | +2.02 |
| **42** | partial cone | 244.30 | +5.80 | 1,2,4-alternate | 339.97 | +4.69 | 1,3,5,7-alternate | 443.67 | +12.08 |
| **43** | 1,3-alternate | 244.31 | +5.81 | 1,2,4-alternate | 339.97 | +4.69 | ¾ cone | 443.69 | +12.10 |
| **44** | partial cone | 244.31 | +5.81 | 1,3,5-alternate | 339.99 | +4.71 | 1,2,4,6,8- alternate | 443.86 | +12.27 |
| **45** | 1,3-alternate | 244.32 | +5.82 | 1,2,4-alternate | 340.02 | +4.74 | 1,2,4,6,8- alternate | 443.87 | +12.28 |
| **46** | partial cone | 244.35 | +5.85 | 1,3,5-alternate | 340.03 | +4.75 | 1,2,4,6,8- alternate | 443.91 | +12.32 |
| **47** | 1,3-alternate | 244.40 | +5.90 | 1,3,5-alternate | 340.05 | +4.77 | 1,2,4,6,8- alternate | 443.96 | +12.37 |
| **48** | partial cone | 244.42 | +5.92 | 1,2,4-alternate | 340.06 | +4.78 | 1,2,4,6,8- alternate | 443.98 | +12.39 |
| **49** | partial cone | 244.43 | +5.93 | 1,2,4-alternate | 340.08 | +4.80 | 1,2,4,6,8- alternate | 444.00 | +12.41 |

**Supplementary Table 2 -** Docking scores of the top 10 docked conformers for 4-sulfonic calix[4]arene, 4-sulfonic calix[6]arene and 4-sulfonic calix[8]arene in AIM2 HIN domain (PDB: 3RN5) using MOE.

| MOE | | | | | |
|---|---|---|---|---|---|
| **4-Sulfonic calix[4]arene** | | **4-Sulfonic calix[6]arene** | | **4-Sulfonic calix[8]arene** | |
| **Docking conformation** | **Docking score (Kcal/mol)** | **Docking conformation** | **Docking score (Kcal/mol)** | **Docking conformation** | **Docking score (Kcal/mol)** |
| 1,2-alternate | -6.7 | 1,3-alternate | -8.4 | 1,2,4,6,8- alternate | -10.2 |
| Cone | -5.8 | 1,3- alternate | -8.4 | 1,2,4,6,8- alternate | -9.9 |
| 1,2-alternate | -5.5 | 1,3- alternate | -8.3 | ¾ cone | -9.9 |
| 1,2-alternate | -5.5 | partial cone | -8.3 | 1,2,4,6,8- alternate | -9.8 |
| 1,2-alternate | -5.5 | partial cone | -8.1 | 1,2,4,6,8- alternate | -9.6 |
| 1,2-alternate | -5.5 | partial cone | -8.1 | ¾ cone | -9.5 |
| 1,2-alternate | -5.5 | 1,3,5- alternate | -8.1 | 1,2-alternate | -9.4 |
| 1,2-alternate | -5.0 | 1,2,4- alternate | -8.1 | 1,2,4,6,8- alternate | -9.4 |
| 1,2-alternate | -4.5 | 1,4- alternate | -7.8 | Pleated loop | -9.4 |
| 1,2-alternate | -4.4 | 1,4- alternate | -7.8 | 1,3,5,7-alternate | -9.3 |

**Supplementary Table 3 -** Docking scores of the top 20 docked conformers for 4-sulfocalix[4]arene, 4-sulfocalix[6]arene and 4-sulfocalix[8]arene in in AIM2 HIN domain (PDB: 3RN5) using Autodock Vina.

| Conformer docking pose | AutoDock Vina | | | | | | | | | | | | | | | | | | | | | | | | | | | |
|---|---|---|---|---|---|---|---|---|---|---|---|---|---|---|---|---|---|---|---|---|---|---|---|---|---|---|---|---|
| | 4-Sulfonic calix[4]arene | | | | 4-Sulfonic calix[6]arene | | | | | | | | 4-Sulfonic calix[8]arene | | | | | | | | | | | | | | | |
| | Docking score (Kcal/mol) | | | | Docking score (Kcal/mol) | | | | | | | | Docking score (Kcal/mol) | | | | | | | | | | | | | | | |
| | 1,2-alternate | partial cone | 1,3-alternate | cone | 1,2-alternate | partial cone | 1,4-alternate | 1,2,4-alternate | 1,2,3-alternate | 1,3-alternate | 1,3,5-alternate | cone | 1,2-alternate | 1,3,5,7-alternate | 1,4-alternate | 1,2,4-alternate | 1,2,4,6,8-alternate | 1,5-alternate | 1,3,5-alternate | 1,4,5-alternate | 1,2,3,4-alternate | 1,3,4,6-alternate | 1,4,5,6-alternate | 1,2,5-alternate | 1,3,4,5-alternate | 1,2,4,5-alternate | ¾ cone | Pleated loop |
| **1** | -16.2 | -16.1 | -15.3 | -16.1 | -16.0 | -16.2 | -15.2 | -16.3 | -16.1 | -15.4 | -15.3 | -15.8 | -13.6 | -15.9 | -14.9 | -15.4 | -14.5 | -15.6 | -15.9 | -14.4 | -15.7 | 14.7 | -15.7 | -13.4 | -16.0 | -15.7 | -14.4 | -14.7 |
| **2** | -15.2 | -16.1 | -15.1 | -15.6 | -14.6 | -16.2 | -14.4 | -15.2 | -15.1 | -14.6 | -14.6 | -15.8 | -13.5 | -15.3 | -14.3 | -14.9 | -14.2 | -15.3 | -15.6 | -14.0 | -14.9 | -13.9 | -14.9 | -13.0 | -13.1 | -14.9 | -13.4 | -14.5 |
| **3** | -15.0 | -14.3 | -14.9 | -13.8 | -14.3 | -16.0 | -14.3 | -14.8 | -14.4 | -13.2 | -14.4 | -15.7 | -13.2 | -14.9 | -14.2 | -14.3 | -14.1 | -13.6 | -14.7 | -13.7 | -14.5 | -13.8 | -14.2 | -12.8 | -12.8 | -14.5 | -13.1 | -13.6 |
| **4** | -14.9 | -14.2 | -14.7 | -13.7 | -14.2 | -14.6 | -14.2 | -14.8 | -14.3 | -13.1 | -13.4 | -15.2 | -13.2 | -13.9 | -14.1 | -14.3 | -13.9 | -13.4 | -14.7 | -13.6 | -14.5 | -13.6 | -14.1 | -12.7 | -12.7 | -14.1 | -13.0 | -13.6 |
| **5** | -14.8 | -13.5 | -14.5 | -13.6 | -14.1 | -14.5 | -14.1 | -14.7 | -14.3 | -12.5 | -13.3 | -15.1 | -13.2 | -13.8 | -13.2 | -14.1 | -13.8 | -13.3 | -14.6 | -13.6 | -14.0 | -13.6 | -14.0 | -12.5 | -12.6 | -13.9 | -12.8 | -13.3 |
| **6** | -14.6 | -13.2 | -14.4 | -13.5 | -13.7 | -13.0 | -13.6 | -14.2 | -14.0 | -12.4 | -13.1 | -14.7 | -13.1 | -13.2 | -13.1 | -13.1 | -13.4 | -12.9 | -14.4 | -13.4 | -13.9 | -13.6 | -13.9 | -12.3 | -12.4 | -13.7 | -12.7 | -13.3 |
| **7** | -14.0 | -13.2 | -14.4 | -13.3 | -13.1 | -13.0 | -13.5 | -14.1 | -13.9 | -12.4 | -13.1 | -13.7 | -12.3 | -13.0 | -13.0 | -12.8 | -13.3 | -12.5 | -14.3 | -13.3 | -13.8 | -13.5 | -12.6 | -12.3 | -12.2 | -13.1 | -12.7 | -13.3 |
| **8** | -13.3 | -13.2 | -14.4 | -13.1 | -12.8 | -12.9 | -13.3 | -13.8 | -13.9 | -12.1 | -13.0 | -13.3 | -12.2 | -13.0 | -12.9 | -12.5 | -13.0 | -12.5 | -13.9 | -13.2 | -13.5 | -13.4 | -12.3 | -11.7 | -11.7 | -12.8 | -12.6 | -13.1 |
| **9** | -13.1 | -13.2 | -14.0 | -12.7 | -12.5 | -12.6 | -13.0 | -13.4 | -13.9 | -12.1 | -12.8 | -13.3 | -12.1 | -12.8 | -12.9 | -12.2 | -11.9 | -12.2 | -13.7 | -13.0 | -13.3 | -13.4 | -12.2 | -11.7 | -11.5 | -12.8 | -12.5 | -12.9 |
| **10** | -13.0 | -13.1 | -13.9 | -12.7 | -12.4 | -12.5 | -12.8 | -13.1 | -13.6 | -12.1 | -12.8 | -13.2 | -11.9 | -12.8 | -12.6 | -12.1 | -11.6 | -12.2 | -13.7 | -13.0 | -13.3 | -13.3 | -12.1 | -11.6 | -11.5 | -12.8 | -12.4 | -12.6 |
| **11** | -12.9 | -12.8 | -13.7 | -12.5 | -12.3 | -12.4 | -12.5 | -12.9 | -13.5 | -12.0 | -12.7 | -13.1 | -11.7 | -12.8 | -12.5 | -12.0 | -11.5 | -12.2 | -12.6 | -12.8 | -13.0 | -13.1 | -11.9 | -11.6 | -11.4 | -12.6 | -11.9 | -12.3 |
| **12** | -12.9 | -12.7 | -13.3 | -12.5 | -12.0 | -12.2 | -12.5 | -12.5 | -13.3 | -11.8 | -12.6 | -12.5 | -11.7 | -12.5 | -12.5 | -11.9 | -11.2 | -12.2 | -12.5 | -12.8 | -12.6 | -13.0 | -11.8 | -11.1 | -11.3 | -12.6 | -11.6 | -11.9 |
| **13** | -12.6 | -12.3 | -13.2 | -12.3 | -11.9 | -12.0 | -12.5 | -12.2 | -13.3 | -11.7 | -12.6 | -12.4 | -11.3 | -12.5 | -12.5 | -11.8 | -11.2 | -12.1 | -12.3 | -12.8 | -12.3 | -12.9 | -11.8 | -11.1 | -11.0 | -12.2 | -11.5 | -11.9 |
| **14** | -12.5 | -12.1 | -12.5 | -12.0 | -11.9 | -11.9 | -12.1 | -12.0 | -13.2 | -11.6 | -12.5 | -12.0 | -11.3 | -12.4 | -12.2 | -11.4 | -10.8 | -12.0 | -12.0 | -12.5 | -12.1 | -12.7 | -11.5 | -10.9 | -10.7 | -12.2 | -11.4 | -11.6 |
| **15** | -12.4 | -12.0 | -12.2 | -11.5 | -11.9 | -11.8 | -12.1 | -11.5 | -13.2 | -11.6 | -12.5 | -11.9 | -11.1 | -12.3 | -12.0 | -11.3 | -10.7 | -12.0 | -11.8 | -12.5 | -11.8 | -12.6 | -11.5 | -10.9 | -10.5 | -11.9 | -11.3 | -11.0 |
| **16** | -12.4 | -12.0 | -12.1 | -11.5 | -11.9 | -11.7 | -12.0 | -11.4 | -13.2 | -11.3 | -12.1 | -11.8 | -11.1 | -12.3 | -12.0 | -11.3 | -10.5 | -11.9 | -11.7 | -12.5 | -11.6 | -12.5 | -11.4 | -10.8 | -10.4 | -11.7 | -10.9 | -10.9 |
| **17** | -12.2 | -11.9 | -12.0 | -11.2 | -11.8 | -11.7 | -11.9 | -11.2 | -13.0 | -11.2 | -12.0 | -11.7 | -10.8 | -12.1 | -12.0 | -11.3 | -10.5 | -11.8 | -11.7 | -12.4 | -11.3 | -12.3 | -11.4 | -10.7 | -10.3 | -11.7 | -10.9 | -10.6 |
| **18** | -12.1 | -11.7 | -12.0 | -11.1 | -11.8 | -11.3 | -11.9 | -11.2 | -12.8 | -11.0 | -11.7 | -11.7 | -10.7 | -12.1 | -11.8 | -11.3 | -10.4 | -11.4 | -11.5 | -12.2 | -11.2 | -12.2 | -11.1 | -10.7 | -10.3 | -11.5 | -10.9 | -10.4 |
| **19** | -11.9 | -11.5 | -12.0 | -11.1 | -11.6 | -11.3 | -11.7 | -11.1 | -12.4 | -11.0 | -11.7 | -11.4 | -10.6 | -11.9 | -11.7 | -11.3 | -10.3 | -11.3 | -11.2 | -12.1 | -10.7 | -12.0 | -11.0 | -10.7 | -10.3 | -11.3 | -10.8 | -10.4 |
| **20** | -11.7 | -11.4 | -11.9 | -10.7 | -11.5 | -11.2 | -11.3 | -10.9 | -12.2 | -10.8 | -11.6 | -11.3 | -10.2 | -11.8 | -11.7 | -11.1 | -10.3 | -11.2 | -11.1 | -12.1 | -10.6 | -11.9 | -11.0 | -10.5 | -10.1 | -11.2 | -10.7 | -10.4 |

Supplementary Figure 5

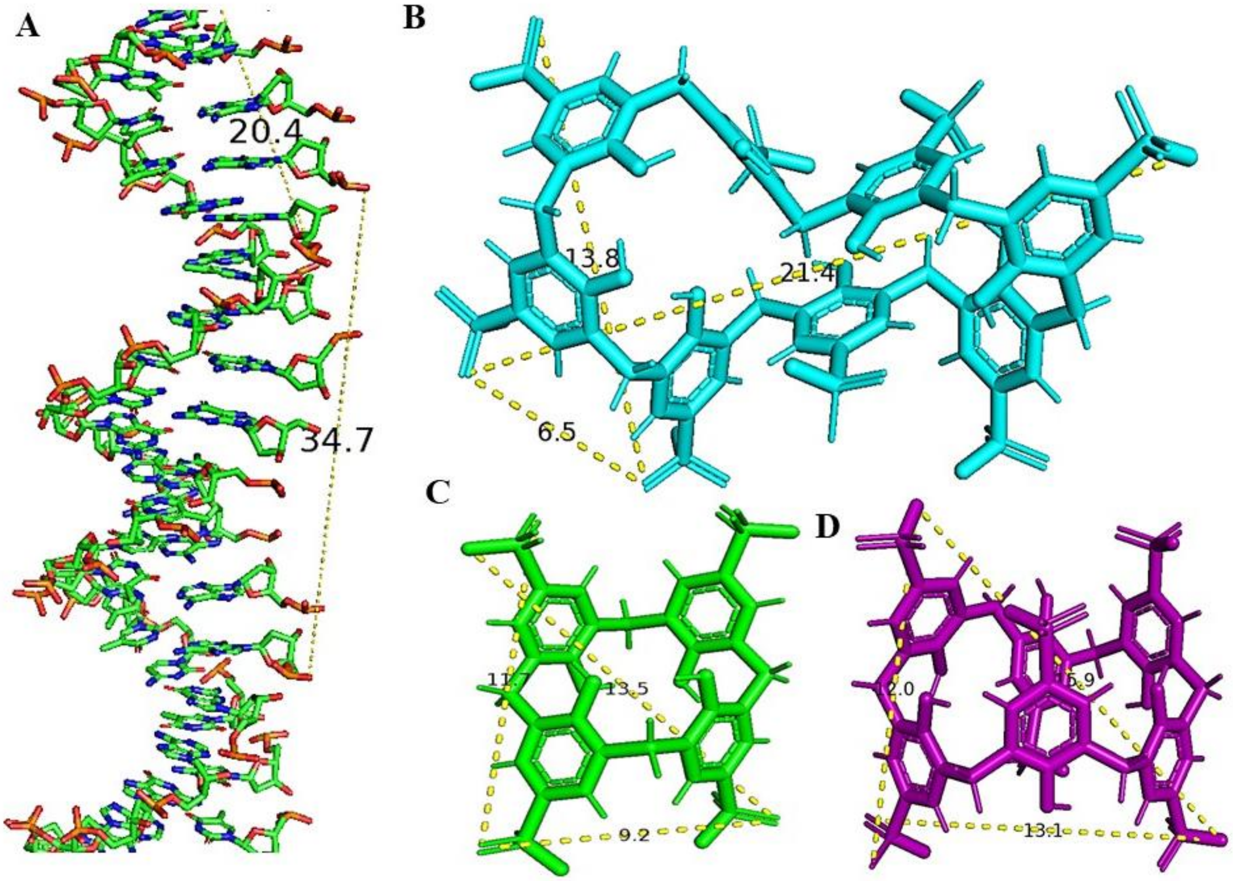


**Supplementary Figure 5 -** Largest measured O-O distances (yellow dashed lines) **(A)** co-crystallized dsDNA in AIM2 for minor groove and major groove regions (PDB: 3RN5), **(B)** 4-sulfonic calix[8]arene, **(C)** 4-sulfonic calix[4]arene, **(D)** 4-sulfonic calix[6]arene. Image created using Pymol.

Supplementary Figure 6

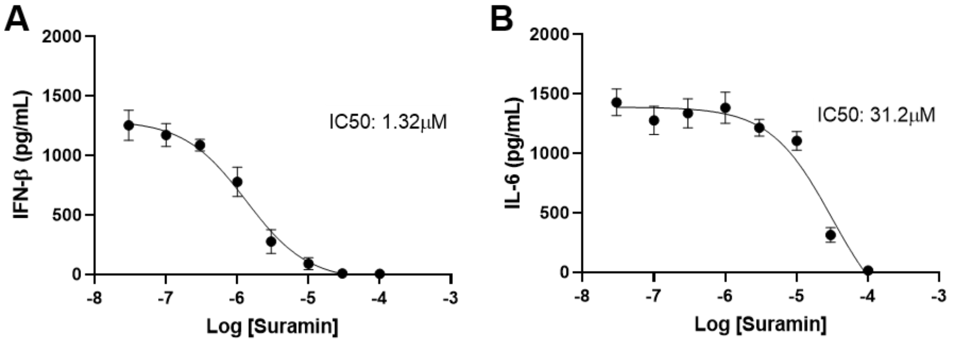


**Supplementary Figure 6 – Suramin is an inhibitor of dsDNA-induced IFN-β release and CpG-induced IL-6 release. (A)** IFN-β release into the supernatants of bone marrow derived macrophages (BMDMs). BMDMs were treated with the indicated concentration of suramin (0.03 – 100 µM) before transfection with poly dA:dT (1 µg $mL^{-1}$, 6 h) (n=4). **(B)** IL-6 release in the supernatants of BMDMs pre-treated with the indicated concentration of suramin (0.03 – 100 µM) and stimulated with CpG DNA (1 µM, 6 h) (n=4). Concentration-response curves were fitted using a four parameter logistical (4PL) model. Values shown are mean ± the SEM.

Supplementary Figure 7

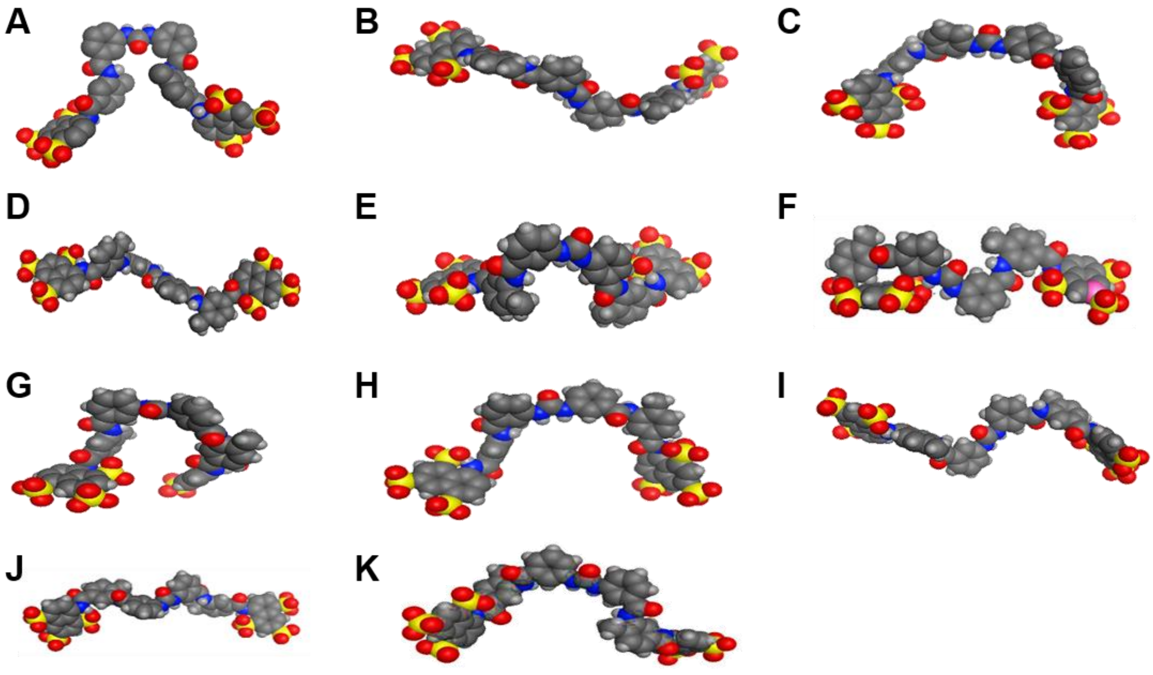


**Supplementary Figure 7 -** Sphere models for the suramin conformers extracted from the crystal structures showing various conformations of suramin including elongated and non-elongated conformers observed having two possible binding modes for some within the same crystal structure **(A)** PDB: 7AH8 **(B)** PDB: 4X3U-1 **(C)** PDB: 4X3U-2 **(D)** PDB: 3GAN-1 **(E)** PDB: 3GAN-2 **(F)** PDB: 6Z7B **(G)** PDB: 6CE2 **(H)** PDB: 4YV5 **(I)** PDB: 2NYR **(J)** PDB: 2H9T **(K)** PDB: 3BJW.

**Supplementary Table 4 -** PDB crystal structures of suramin with their protein targets and therapeutic application with their reported $IC_{50}$/Ki/$\Delta$G values and calculated potential energies of the suramin conformers**.**

| Suramin Conformer no. | Potential energy of the conformer (Kcal/mol) | Crystal structure resolution (Å) | R factor | Name of protein | Therapeutic area | $IC_{50}$/Ki/$\Delta$G |
|---|---|---|---|---|---|---|
| **PDB: 7AH8 (*1*)** | 719.78 | 2.7 | 0.22 | NF-Y | Anti-cancer | $\Delta$G = −12.1 kcal/mol |
| **PDB:**<br>**4X3U-1 (*2*)**<br>**4X3U-2 (*2*)** | 1052.38<br>1419.18 | 1.6 | 0.16 | Chromobox homolog 7 (CBX7) chromodomain | Senescence control, cancer prevention, and stem cell lineage specification | $IC_{50}$ = 8.1 µM |
| **PDB:**<br>**3GAN-1**<br>**3GAN-2** | 866.01<br>1662.92 | 2.0 | 0.19 | Arabidopsis Thaliana AT3G22680 | Anti-parasitic | - |
| **PDB: 6Z7B (*3*)** | 648.86 | 1.8 | 0.19 | Variant Surface Glycoprotein VSGsur | Early-stage treatment for African trypanosomiasis | $IC_{50}$ = 8.4 µM |
| **PDB: 6CE2 (*4*)** | 623.55 | 2.1 | 0.22 | Myotoxin I (MjTX-I) from Bothrops moojeni | Adjuvant of conventional serum therapy against local myonecrosis as a neutralizing agent | Ki = <1 µM |
| **PDB: 4YV5 (*5*)** | 744.65 | 1.9 | 0.19 | Myotoxin II from snake Bothrops moojeni | Adjuvant of conventional serum therapy against local myonecrosis as a neutralizing agent for snake venum | $IC_{50}$= 5.8 µM |
| **PDB: 2NYR (*6*)** | 523.84 | 2.0 | 0.20 | Human $NAD^+$ - Dependent Deacetylase SIRT5 | Treatment of human metabolic and neurological diseases and cancer | $IC_{50}$= 22 µM |
| **PDB: 2H9T (*7*)** | 618.17 | 2.4 | 0.18 | Human Alpha-Thrombin | Anti-thrombin | Ki = <1 µM |
| **PDB: 3BJW (*8*)** | 489.12 | 2.3 | 0.21 | Ecarpholin S | Neutralizing agent | $\Delta$G = -19.8 kcal/mol (protein dimer with two suramin)<br>$\Delta$G = -28.2 kcal/mol. (trimer, one suramin/subunit) |

**Supplementary Table 5 -** Suramin conformers were docked rigidly in the HIN domain (monomer) and between the 2 HIN domains (dimer) (PDB: 3RN5) using Autodock Vina. The maximum measured O-O distance and docking scores of the cocrystal structures of suramin for the top 20 docked poses.

| | Docking scores (Kcal/mol) | | | | | | | | | | |
|---|---|---|---|---|---|---|---|---|---|---|---|
| **PDB** | **1** | **2** | **3** | **4** | **5** | **6** | **7** | **8** | **9** | **10** | **O-O distance (Å)** |
| **7AH8(*1*) Monomer** | -21.9 | -21.4 | -21.3 | -20.5 | -20.4 | -20.2 | -20.2 | -20.1 | -20.1 | -20.1 | 27.1 |
| **Dimer** | -22.4 | -20.8 | -20.7 | -20.5 | -19.9 | -19.9 | -19.5 | -18.5 | -18.5 | -17.4 | |
| **4X3U_1 (*2*) Monomer** | -20.3 | -20.1 | -19.9 | -19.1 | -18.6 | -18.6 | -18.5 | -18.4 | -18.1 | -18.1 | 29.8 |
| **Dimer** | -21.6 | -21.2 | -20.2 | -20.2 | -20.0 | -19.8 | -19.5 | -19.4 | -19.2 | -19.1 | |
| **4X3U_2 (*2*) Monomer** | -24.6 | -23.0 | -21.8 | -20.9 | -20.7 | -20.1 | -19.9 | -19.8 | -19.7 | -19.5 | 27.4 |
| **Dimer** | -22.5 | -20.9 | -20.6 | -20.1 | -20.0 | -19.8 | -19.6 | -19.1 | -18.7 | -18.4 | |
| **3GAN_1 Monomer** | -24.5 | -23.0 | -22.6 | -21.9 | -21.4 | -21.1 | -21.0 | -19.5 | -19.5 | -19.2 | 30.8 |
| **Dimer** | -23.0 | -21.6 | -21.5 | -20.5 | -20.4 | -20.3 | -20.3 | -20.2 | -20.2 | -20.1 | |
| **3GAN_2 Monomer** | -24.5 | -23.0 | -22.7 | -21.7 | -21.5 | -20.8 | -20.7 | -20.6 | -20.4 | -19.6 | 30.8 |
| **Dimer** | -23.0 | -21.7 | -20.5 | -20.4 | -20.2 | -20.2 | -20.2 | -19.9 | -19.9 | -19.8 | |
| **6Z7B(*3*) Monomer** | -24.3 | -22.0 | -21.4 | -21.3 | -21.2 | -20.0 | -19.7 | -19.5 | -19.5 | -19.4 | 28.3 |
| **Dimer** | -22.1 | -21.4 | -21.3 | -20.8 | -20.2 | -20.2 | -19.1 | -18.8 | -18.8 | -18.7 | |
| **6CE2(*4*) Monomer** | -19.6 | -19.6 | -19.1 | -19.1 | -19.1 | -19.0 | -18.6 | -18.6 | -18.5 | -19.8 | 20.7 |
| **Dimer** | -20.1 | -19.2 | -19.0 | -18.6 | -18.0 | -18.0 | -18.0 | -17.8 | -17.8 | -17.8 | |
| **4YV5(*5*) Monomer** | -27.3 | -26.1 | -24.9 | -24.8 | -24.3 | -24.0 | -23.8 | -23.7 | -23.2 | -22.2 | 29.5 |
| **Dimer** | -24.9 | -24.3 | -23.3 | -21.8 | -21.8 | -20.8 | -20.6 | -20.6 | -19.8 | -19.8 | |
| **2NYR(*6*) Monomer** | -23.3 | -22.8 | -22.8 | -22.0 | -21.7 | -21.4 | -20.7 | -19.6 | -19.5 | -19.4 | 34.6 |
| **Dimer** | -22.8 | -22.7 | -22.7 | -22.0 | -22.0 | -21.2 | -20.6 | -20.4 | -19.8 | -19.6 | |
| **2H9T(*7*)** | -23.9 | -23.8 | -23.5 | -22.7 | -22.3 | -21.4 | -20.1 | -19.7 | -19.6 | -19.4 | 35.3 |

| | | | | | | | | | | | |
|---|---|---|---|---|---|---|---|---|---|---|---|
| **Monomer** | | | | | | | | | | | |
| **Dimer** | -23.5 | -23.2 | -22.3 | -21.8 | -20.3 | -19.2 | -19.0 | -18.8 | -18.3 | -17.9 | |
| **3BJW(*8*)**<br>**Monomer** | -26.3 | -24.5 | -23.7 | -22.0 | -21.3 | -21.2 | -21.0 | -20.6 | -20.4 | -20.3 | 36.7 |
| **Dimer** | -24.5 | -21.9 | -21.1 | -20.7 | -20.5 | -20.2 | -20.0 | -19.9 | -19.6 | -19.4 | |

**Supplementary Table 6 -** Different energy minimized conformers of suramin in their ionized forms at pH 7.4 after a conformational search in MOE with calculated potential energies.

| Conformer no. | Potential energy (Kcal/mol) | Urea Conformation | Conformer no. | Potential energy (Kcal/mol) | Urea Conformation |
|---|---|---|---|---|---|
| **1** | 62.07 | trans/trans | **15** | 253.07 | trans/trans |
| **2** | 162.52 | trans/trans | **16** | 253.25 | trans/trans |
| **3** | 163.48 | trans/trans | **17** | 254.23 | trans/cis |
| **4** | 165.69 | trans/trans | **18** | 256.18 | trans/trans |
| **5** | 166.08 | trans/trans | **19** | 257.59 | trans/trans |
| **6** | 173.29 | trans/trans | **20** | 263.01 | trans/trans |
| **7** | 186.32 | trans/trans | **21** | 267.36 | trans/trans |
| **8** | 212.21 | trans/trans | **22** | 269.22 | trans/cis |
| **9** | 247.18 | trans/trans | **23** | 270.06 | trans/cis |
| **10** | 248.87 | trans/cis | **24** | 271.92 | trans/trans |
| **11** | 249.48 | trans/trans | **25** | 274.76 | trans/cis |
| **12** | 250.56 | trans/trans | **26** | 285.96 | trans/cis |
| **13** | 251.00 | trans/trans | **27** | 291.35 | trans/trans |
| **14** | 251.21 | trans/trans | **28** | 317.64 | cis/cis |

**Supplementary Table 7 -** Docking of suramin conformers in the AIM2 HIN domain (monomer) (PDB: 3RN5). Docking scores for the energy minimized forms of suramin (obtained from conformational search in MOE) using Autodock Vina through rigid docking.

| | **Conformer no.** | | | | | | | | | |
|---|---|---|---|---|---|---|---|---|---|---|
| **Docking score (Kcal/mol)** | **1** | **2** | **3** | **4** | **5** | **6** | **7** | **8** | **9** | **10** |
| | -21.2 | -23.5 | -23.4 | -23.5 | -20.8 | -22.1 | -25.2 | -24.9 | -22.3 | -23.1 |
| | -20.0 | -21.5 | -21.9 | -21.7 | -20.2 | -22.0 | -22.2 | -21.6 | -21.8 | -21.7 |
| | -19.8 | -21.3 | -20.1 | -21.5 | -20.2 | -21.9 | -21.3 | -21.5 | -20.0 | -20.4 |
| | -19.7 | -21.2 | -20.0 | -20.4 | -19.7 | -21.7 | -21.3 | -20.8 | -19.9 | -20.4 |
| | -19.7 | -20.9 | -19.9 | -19.9 | -19.5 | -21.6 | -21.2 | -19.8 | -19.8 | -19.6 |
| | -19.6 | -20.6 | -19.5 | -19.3 | -19.2 | -20.3 | -21.1 | -19.4 | -19.5 | -19.5 |
| | -19.4 | -20.0 | -19.0 | -19.2 | -19.1 | -20.0 | -21.0 | -19.0 | -19.2 | -19.4 |
| | -19.2 | -19.9 | -18.9 | -19.1 | -19.0 | -19.7 | -20.7 | - | -19.2 | -19.3 |
| | -19.1 | -19.8 | -18.8 | -19.0 | -18.1 | -19.6 | -20.4 | - | -19.0 | -19.1 |

| | **Conformer no.** | | | | | | | | | | |
|---|---|---|---|---|---|---|---|---|---|---|---|
| **Docking score (Kcal/mol)** | **11** | **12** | **13** | **14** | **15** | **16** | **17** | **18** | **19** | **20** | **21** |
| | -20.1 | -20.6 | -22.0 | -22.1 | -23.0 | -24.7 | -23.6 | -24.9 | -22.0 | -23.7 | -21.7 |
| | -19.9 | -19.5 | -21.9 | -21.3 | -22.8 | -22.2 | -22.9 | -23.8 | -21.2 | -23.6 | -21.2 |
| | -19.5 | -19.2 | -21.0 | -21.2 | -21.0 | -21.7 | -20.5 | -22.7 | -20.5 | -23.2 | -19.9 |
| | -19.2 | -19.1 | -20.9 | -20.9 | -20.6 | -21.1 | -20.5 | -21.1 | -20.0 | -22.2 | -19.8 |
| | -19.0 | -18.6 | -20.9 | -20.0 | -20.0 | -21.0 | -20.3 | -20.6 | -19.9 | -22.0 | -19.7 |
| | -19.0 | -18.2 | -20.6 | -19.7 | -19.8 | -20.7 | -19.9 | -20.2 | -19.6 | -21.7 | -19.7 |
| | -18.9 | -18.0 | -20.0 | -19.7 | -19.5 | -20.6 | -19.7 | -20.1 | -19.2 | -21.5 | -19.5 |
| | -18.8 | -17.7 | -19.9 | -19.6 | -18.7 | -20.4 | -19.2 | -20.1 | -19.1 | -20.9 | -19.3 |
| | -18.7 | -17.5 | -19.6 | -19.2 | -18.4 | -19.8 | -19.0 | -19.9 | -19.0 | -20.7 | -19.3 |

| | **Conformer no.** | | | | | | | |
|---|---|---|---|---|---|---|---|---|
| **Docking score (Kcal/mol)** | **22** | **23** | **24** | **25** | **26** | **27** | **28** | **29** |
| | -21.7 | -22.5 | -22.4 | -22.2 | -23.9 | -20.8 | -22.9 | -21.0 |
| | -20.9 | -22.3 | -21.2 | -21.0 | -23.8 | -20.6 | -21.7 | -20.7 |
| | -20.4 | -21.0 | -20.4 | -20.9 | -23.5 | -19.9 | -20.5 | -20.6 |
| | -19.9 | -20.9 | -20.1 | -20.4 | -22.7 | -19.8 | -20.5 | -20.5 |
| | -19.9 | -20.8 | -20.0 | -20.4 | -22.3 | -19.8 | -20.4 | -20.1 |
| | -19.8 | -20.2 | -19.9 | -19.8 | -21.4 | -19.8 | -20.3 | -20.1 |
| | -19.7 | -20.1 | -19.8 | -19.7 | -20.1 | -19.7 | -20.2 | -20.1 |
| | -19.3 | -20.0 | -19.5 | -19.6 | -19.7 | -19.6 | -19.8 | -20.0 |
| | -19.3 | -19.8 | -19.2 | -18.9 | -19.6 | -19.5 | -19.6 | -20.0 |

Supplementary Figure 8

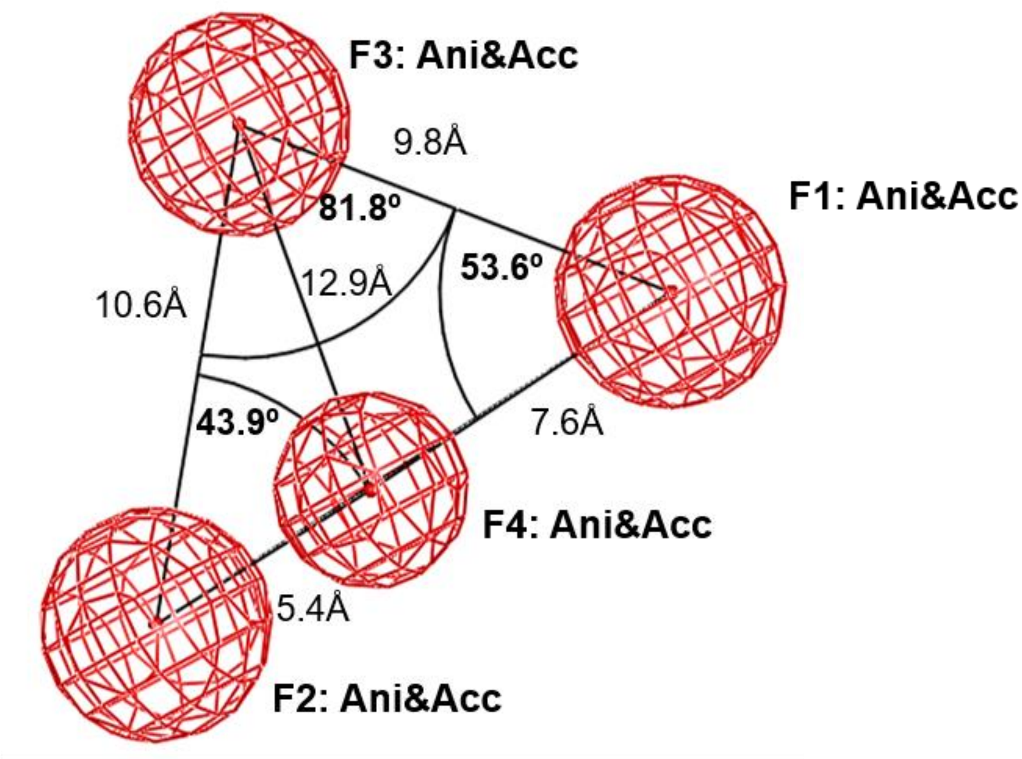


**Supplementary Figure 8 -** Pharmacophore model, derived from the best binding modes from the docking of 4-sulfonic calixarenes in the AIM2 HIN domain (PDB:3RN5) using Autodock Vina and MOE after overlay of the docked poses. The pharmacophore consists of four ionic interaction sites with radii of 2.1Å, 2.4Å, 2.5Å and 2.5Å (red spheres). Distances and dihedral angles are shown.

**Supplementary Table 8 -** Ionic interactions and docking scores of the conformers (conf1 & conf2) for suramin in AIM2 (PDB: 3RN5) using MOE and Autodock Vina. Distances from the sulfonate oxygens of suramin to the positively charged amino acid side chains in the HIN domain binding pocket.

| | MOE (Å) | AutoDock Vina (Å) | |
|---|---|---|---|
| **Interacting residue** | **Suramin (conf1)** | **Suramin (conf1)** | **Suramin (conf2)** |
| **Docking score (Kcal/mol)** | -12.2 | -18.9 | -22.0 |
| **Lys160** | 2.7 | 2.9 | - |
| **Lys162** | 2.9 | 3.1 | - |
| **Lys163** | 2.1 | 2.9 | 2.6 |
| **Lys198** | 3.2 | 3.5 | 3.4 |
| **Lys204** | 2.8 | 2.1 | - |
| **Lys251** | - | - | 3.2 |
| **Arg311** | 1.7* | 2.3* | - |
| **Lys335** | - | - | - |

Supplementary Figure 9

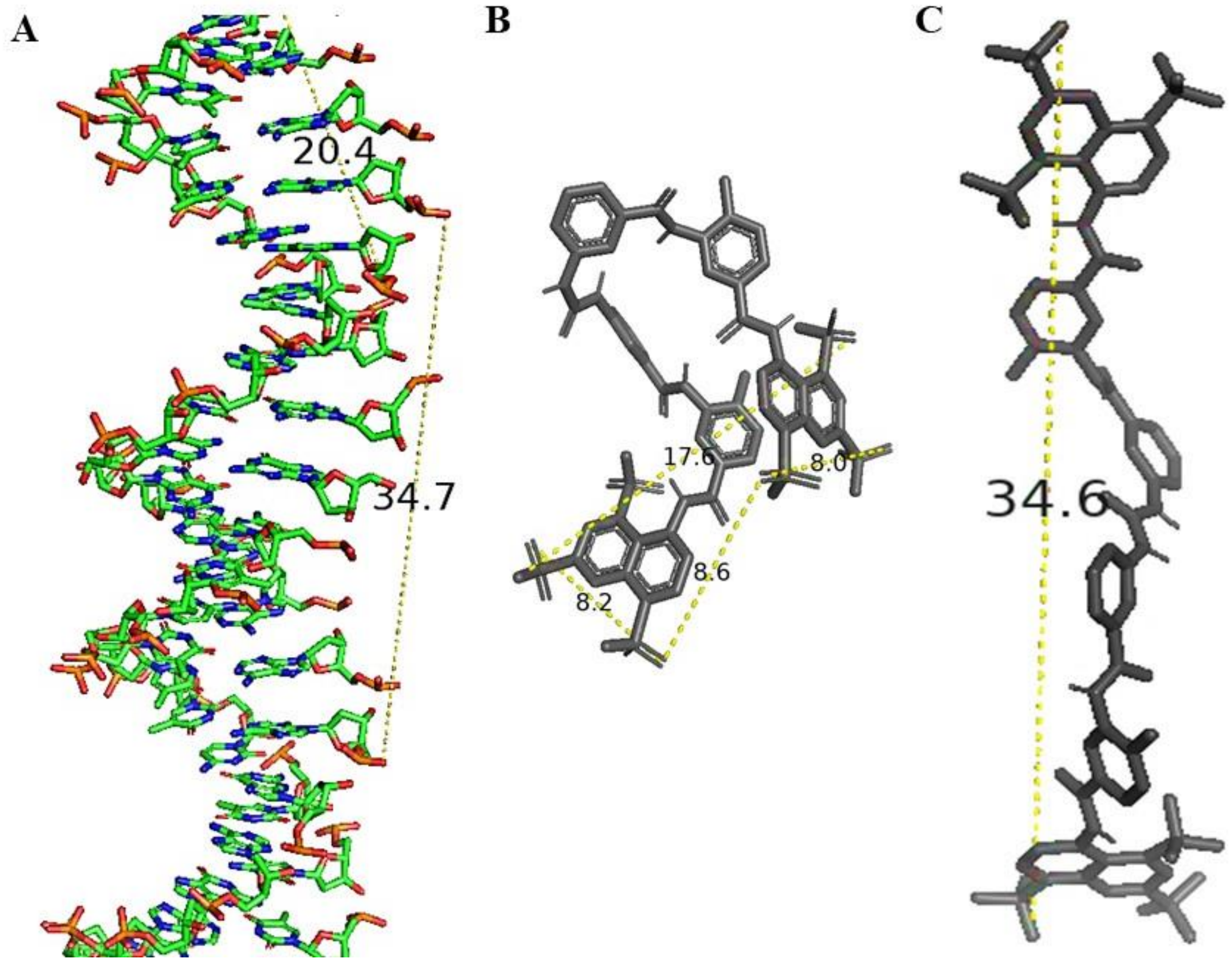


**Supplementary Figure 9 -** Largest measured O-O distances (yellow dashed lines) **(A)** co-crystallized dsDNA in AIM2 for minor groove and major groove regions (PDB: 3RN5)**, (B)** preferred docked pose of suramin in the HIN domain (PDB:3RN5) **(C)** extended conformer of suramin for the preferred pose between the 2 HIN domains of AIM2. Image created using Pymol.

Supplementary Figure 10

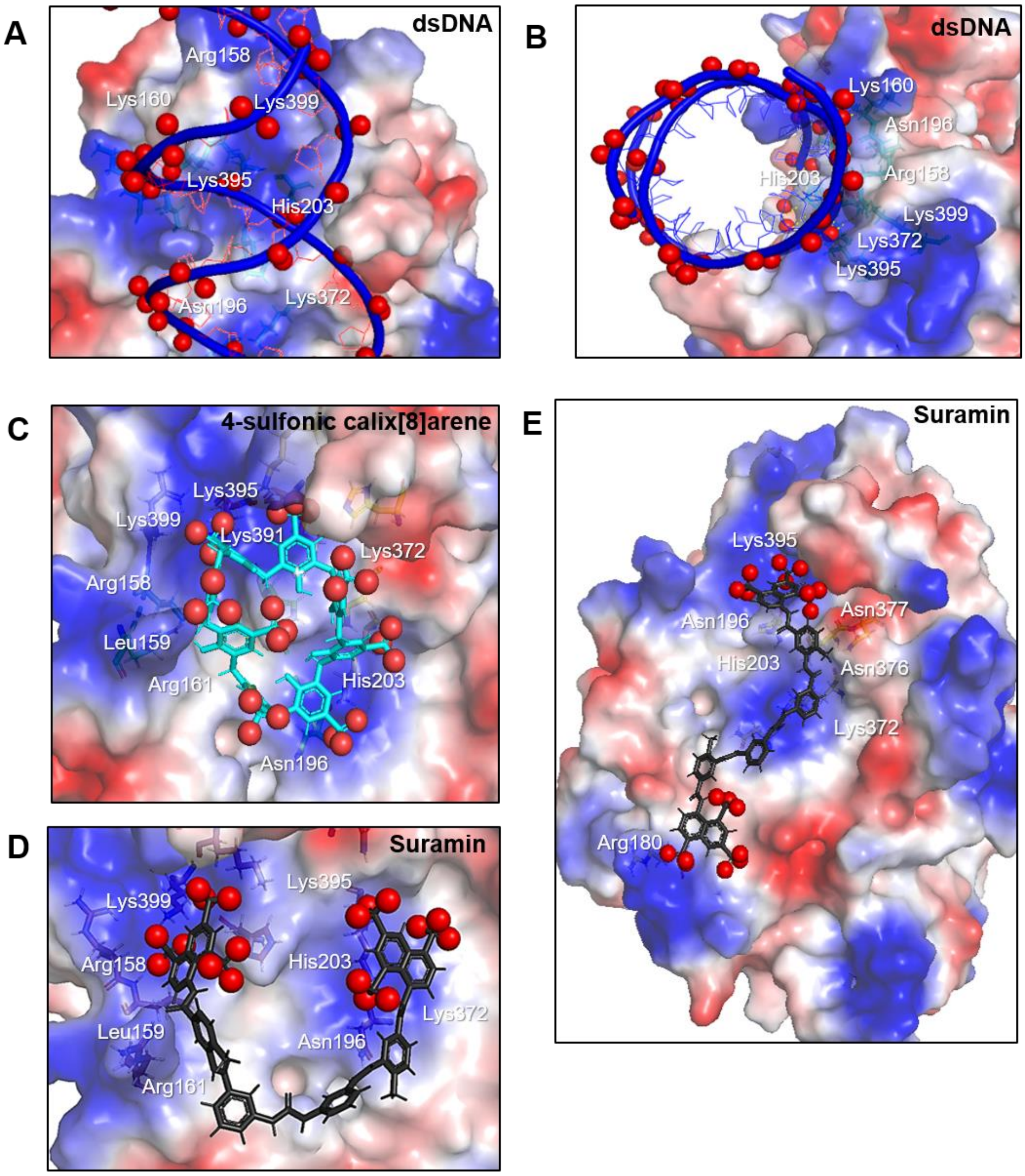


**Supplementary Figure 10 - Modelling of 4-sulfonic calix[8]arene and suramin on the X-ray crystal structure of dsDNA bound cGAS.** Docking validation in mouse cGAS (PDB: 4LEZ) **(A,B)** electrostatic potential map for docked dsDNA (blue) showing interacting residues, **(A)** side view **(B)** top view. For electrostatic potential maps: blue: positive region, red: negative region, white: neutral region, red spheres: negatively charged atoms for the phosphates of dsDNA. Electrostatic potential maps for docking of inhibitors in mouse cGAS (monomer) (PDB: 4LEZ), **(C)** 4-sulfonic calix[8]arene using MOE, **(D)** suramin (conf3) in cGAS (monomer) using Autodock Vina, **(E)** entire cGAS monomer showing the alternate binding mode (extended form) of suramin (grey) (conf4) using MOE mimicking the dsDNA turn with similar interacting amino acid residues seen for the overlaid sulfonates of suramin. For electrostatic potential maps: blue: positive region, red: negative region, white: neutral region, red spheres: negatively charged atoms for the sulfonates of 4-sulfonic calix[8]arene & suramin. Images created using Pymol.

**Supplementary Table 9 -** Docking of 4-sulfonic calix[8]arene and suramin (conf3 & conf4) in cGAS (PDB: 4LEZ) using MOE and Autodock Vina. Docking scores and distances for the ionic interactions of the best docked conformers for 4-sulfonic calix[8]arene and suramin. Distances from the sulfonate oxygen to the positively charged nitrogen atom in the cGAS binding pocket, in comparison with the negatively charge phosphate oxygen of the co-crystallized DNA.

*Sulfonate oxygens forming hydrogen bond with amino acid backbone NH.

| - | | MOE (Å) | | Autodock Vina (Å) | |
|---|---|---|---|---|---|
| - | Co-crystallized DNA | 4-Sulfonic calix[8]arene | Suramin conf3 | | Suramin conf4 |
| **Docking score (Kcal/mol)** | - | -9.5 | -10.9 | -11.2 | -10.6 |
| **Arg158** | 3.5 | 3.2 | 3.2* | 3.6 | - |
| **Leu159** | 3.9* | 2.1* | 2.0* | 2.3* | - |
| **Lys160** | 3.5 | - | - | - | - |
| **Arg161** | - | 3.1 | 2.6* | 3.8* | - |
| **Arg180** | 3.4 | - | - | - | 3.1 |
| **Asn196** | 2.9* | 3.6 | 1.9* | 2.0* | 3.7 |
| **His203** | 3.9 | 3.0 | 3.9 | 3.3 | 3.9 |
| **Lys372** | 2.7 | 3.3 | 3.0 | 3.0 | 3.1* |
| **Asn376** | - | 3.1 | 3.7 | 3.1 | 3.1* |
| **Asn377** | - | - | - | - | 3.4 |
| **Lys391** | - | 2.2* | 3.7 | - | - |
| **Lys395** | 3.1 | 3.4 | 3.2 | 3.4 | 3.0 |
| **Lys399** | 3.9 | 3.0 | 3.0 | 3.4 | - |
| **O-O distance (Å)** | 22.0 | 20.8 | 21.9 | 21.9 | 33.7 |
